# Extracellular matrix mediated stresses spatially bias myoblast fusion and myotube growth

**DOI:** 10.1101/2024.11.22.624831

**Authors:** Yoann Le Toquin, Sushil Dubey, Aleksandra Ardaševa, Lakshmi Balasubramaniam, Emilie Delaune, Valérie Morin, Amin Doostmohammadi, Christophe Marcelle, Benoît Ladoux

## Abstract

Myoblast fusion into multinucleated myotubes is essential for skeletal muscle development and repair, yet how tissue-scale mechanics contributes to this process remains poorly understood. Here, we show that primary myoblasts behave as an evolving active nematic system in which actomyosin-dependent stresses, extracellular-matrix (ECM) remodeling and fusion-driven myotube growth are dynamically coupled. As myoblasts fuse into elongated myotubes, orientational order increases and the nematic field is progressively reshaped. We identify a strong coupling between cellular and ECM nematic organization, whereby cytoskeleton-dependent ECM remodeling stabilizes topological defects and reinforces their associated stress patterns. Fusion events preferentially accumulate near comet-shaped +1/2 defects, which correspond to regions of high compressive stress predicted by our theoretical model. Our findings support a model in which the intrinsic fusion machinery provides fusion competence, while ECM-stabilized nematic stress patterns spatially bias the localization of fusion events. Fusion-driven myotube growth then feeds back on the mechanical landscape, increasing nematogen length and stress magnitude. Together, these results reveal a self-reinforcing biomechanical mechanism that contributes to the organization of myoblast fusion and myotube growth, with potential relevance for developmental and regenerative morphogenesis.

## Introduction

Myoblast fusion is a critical, late-stage event in myogenesis, driving the formation of multinucleated myofibers. This complex process requires precise coordination of cell migration, cell-cell recognition, cytoskeletal remodeling, and membrane fusion^1–3^. Recent breakthroughs have shed light on the molecular players essential for myoblast fusion, alongside newly identified inhibitors that modulate the process^1–5^. *In vitro*, myoblasts generate myofibers that self-organize into striking, aligned patterns, forming contractile bundles within days^6^. Although tension has been shown to be required for sarcomeric assembly in myofibers^6^, the mechanical framework leading to myotube formation and patterning, which heavily relies on myoblast fusion, remains largely unexplored. Unlocking these mechanical principles is critical, as force transduction between cells is known to affect processes such as migration, alignment, or differentiation^7,8^. Understanding the forces at play during muscle fusion might offer insights into fundamental processes that drive tissue architecture and function across the body^9^.

In recent years, there has been a notable increase in the application of soft matter physics to the study of biological systems^10,11^. Various cell types surprisingly exhibit self-organizing behaviors resembling active nematic liquid crystals, through the formation of supracellular domains with local long-range orientational order (nematic order), where cells are aligned to one another, just as rod-shaped particles of liquid crystals do^12^. Interestingly, similar to rod-shaped particles, ordering of cells is perturbed by singularities in their orientational field, called topological defects. These defects have been shown to play a major role in the mechanical regulation of various biological processes including cell death^13^, monolayer rupture^14^, morphogenesis^15^ and cell differentiation^16^. The process of myoblast fusion is intrinsically different from other cell systems, as the biological objects evolve during fusion, myoblasts gradually generating myofibers. It is therefore unknown how cellular ordering and associated self-organization processes evolve during the myoblast to myofiber transition.

Here, we demonstrate that topological defects in the active nematic texture of the myoblast assemblies contribute to shaping the spatial regulation of myoblast fusion *in vitro*. In contrast to previous studies, our system reveals that dynamically evolving material properties are central to muscle-cell differentiation, arising both from the morphological and mechanical changes that cells undergo as they fuse into myotubes and from a previously unrecognized coupling between actomyosin-dependent cellular and extracellular-matrix (ECM) nematic order. As differentiation proceeds, this coupling stabilizes and transmits the orientational field, giving rise to stationary and stable topological defects accompanied by a progressively reinforced mechanical stress landscape which influences the distribution of fusion events in space. Indeed, we find that comet-shaped topological defects act as organizing centers where compressive stresses accumulate, with both the size and magnitude of these stresses increasing over time and strongly correlating with the spatial localization of fusion events. To explain these dynamics, we developed a continuum active nematic model coupled to a self-generated, evolving ECM, which captures the observed increase in enhanced defects-associated stresses throughout the myoblast-to-myotube transition. We then confirmed the model’s predictions by culturing primary myoblasts over a pre-patterned decellularized ECM which allowed total recapitulation of the stable topological defects and associated stresses observed at the end of the previous culture as well as of the predicted spatial bias of fusion locations toward the compressive stress regions associated to the reformed comet-shaped defects. Our findings support a model in which dynamic mechanical interaction between cells and the remodeling of the underlying matrix promote the spatial coordination and amplification of fusion during myotube formation.

## Results

### Primary myoblasts behave as contractile active nematic liquid crystals

We characterized the behavior of primary embryonic muscle progenitors during their transition from myoblasts to myotubes *in vitro*. Primary muscle progenitors were isolated from the hindlimbs of 11-days-old chicken embryos^17^ and cultured for up to 72 hours on a soft deformable Polydimethylsiloxane (PDMS) substrate with stiffness (15 *k*Pa) typical of normal muscle tissues^18,19^. During this period, muscle progenitors undergo differentiation^20–22^, accompanied by massive fusion, leading to the formation of myotubes containing numerous nuclei (Supplementary Video 1).

As primary myoblasts reached confluency, phase-contrast bright field microscopy revealed distinctive patterns of collective self-organization, characterized by the formation of domains where cells adopted similar orientations, typical of nematic systems^23–25^ (Fig. 1a). Over time, the nematic domains of cells displaying similar orientations, recognized by identical colors, dramatically expanded (Fig. 1a). To quantify this organization, we determined the cell population orientation field and calculated the degree of orientational order, defined by the scalar nematic order parameter (*S*) at each time point during the culture period (Fig. 1b, see Methods section). The order parameter increased gradually over time, with its curve slowing down towards the end of the culture period (Fig. 1b), suggesting increased orientational order across the entire cell populations (Extended Data Fig. 1a-b). Simultaneously, through calculation of the winding number parameter, which evaluates the local rotation of the orientation field, we observed the formation of comet- (+1/2) and trefoil-like (-1/2) topological defects, characteristic of a nematic configuration at the intersections of cell domains^12^ (Fig.1c). By analyzing the flow field (i.e. global displacements within consecutive images over time) around +1/2 comet-shaped defects using particle image velocimetry (PIV), we found that flows were oriented towards the tail of the +1/2 defects – a characteristic feature of active contractile systems^26^ (Fig. 1d, Extended Data Fig. 1c). As a consequence of this behavior, we observed a slow displacement of +1/2 defects in the direction of their tail over the culture time (Fig. 1e). Interestingly, this resulted in the merging of topological defects with opposite charges (+1/2 merging with -1/2) and in their annihilation (Fig. 1e-f, Supplementary Video 2). During the myoblast to myotube transition, annihilation events dominated over nucleation of new defects. This led to the global reduction in defect density and to a gradual increase in the distance between defects, which was accompanied by the cellular nematic order increase (Fig. 1b, 1f, Extended Data Fig. 1a-b, d).

**Fig. 1:**
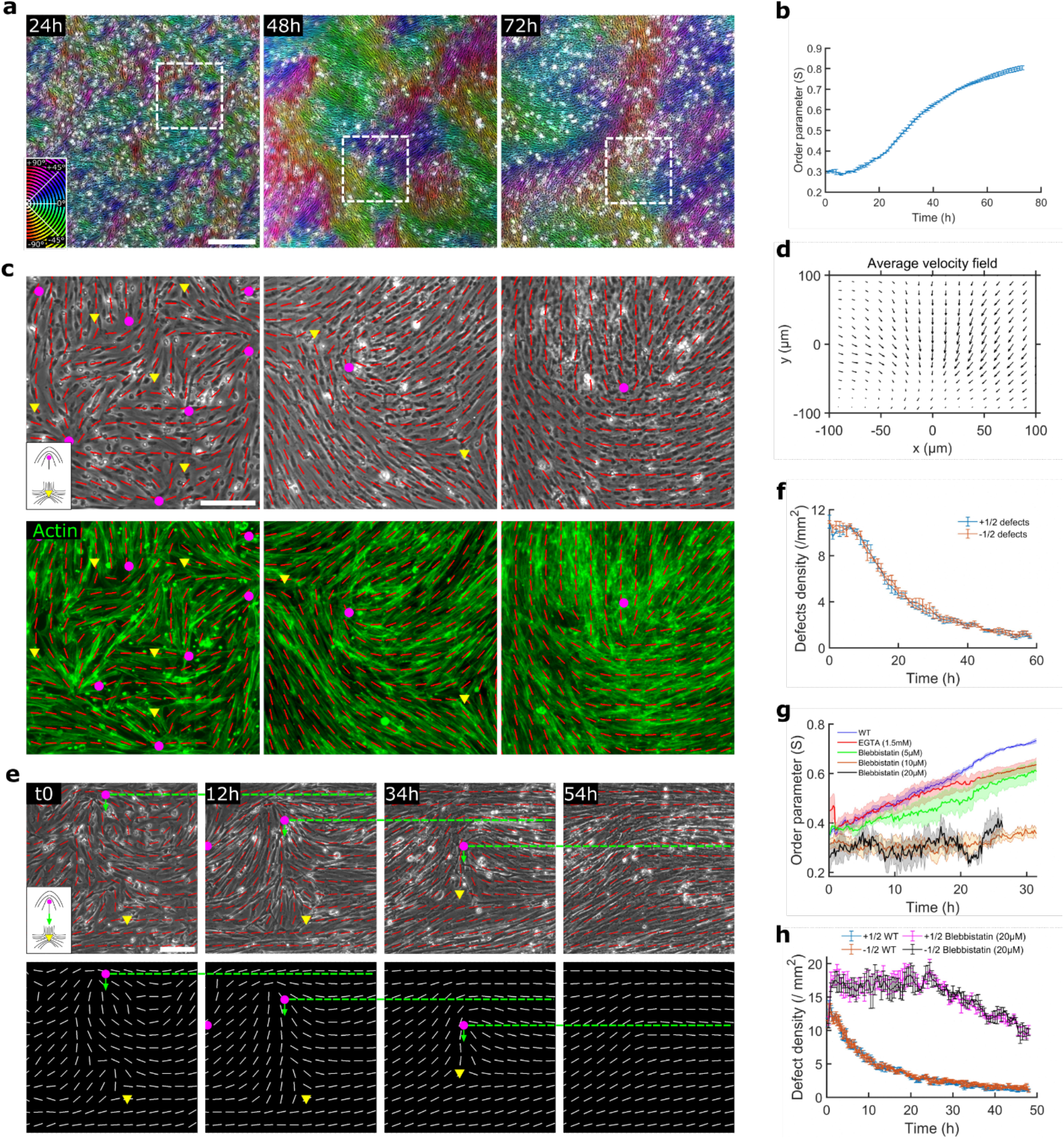
Contractile active nematic behavior of primary myoblasts is dependent on cytoskeletal activity. **a**, Bright-field pseudo-colored according to local orientation of myoblasts cultures fixed at various time-points. Inset shows the color wheel applied to the local orientation of the cells. Scale bar, 500 µm. **b**, Time evolution of the order parameter in primary myoblasts cultures (8×8 nematic directors approx. 200 µm x 200 µm window size, n = 3 independent movies, error bars are standard error (S.E.) from the mean). **c**, Close up showing the boxed region of the corresponding images in (**a**) with phase contrast (top row) or phalloidin (bottom row). All images are overlaid with the averaged local orientation of the cells (red lines) and markings for comet-shape +1/2 (magenta dots) and trefoil-shape -1/2 (yellow triangles) defects. Inset shows a schematic of the shape of the defects. Scale bar, 100 µm. **d**, Average velocity field around +1/2 defects (n=7503 defects from n=3 independent movies). **e**, Time lapse of a motile +1/2 defect from birth to annihilation. Top panels: bright-field overlaid with averaged local orientation of the cells (red lines). Bottom panels: averaged local orientation of the cells (white lines). Dashed lines and green arrows allow to better observe the motion of the +1/2 defect (magenta dots) toward a -1/2 defect (yellow triangle) with whom it fuses. Inset shows a schematic of the +1/2 defect movement towards -1/2 defect. Scale bar, 100 µm. **f**, Time evolution of the defects density in primary myoblasts cultures (n = 3, independent movies, error bars are S.E. from the mean). **g-h**, Time evolution of the order parameter (**g**) and defects density (**h**) in primary myoblasts cultures for control (Dimethyl sulfoxide-DMSO) or drugs (blebbistatin (5,10, 20 µM) and EGTA (1.5mM)) treated cells (averaged over 3 independent movies, error bars are S.E. from the mean).

Importantly, the orientation field obtained from monitoring cell axes through phase-contrast microscopy matched subcellular scale orientation of actin stress fibers (Fig. 1c). As we observed a close correspondence between the global orientation of the cell population and the orientation field of subcellular cytoskeletal (actin) organization (Fig. 1c), we hypothesized that cytoskeletal organization and activity constitute key elements in the evolution of the nematic behavior of the myoblast population. As the myogenic population continued to evolve over time, the process of differentiation and fusion leads to the formation of elongated myotubes, which in turn impacts cytoskeletal activity and shape^27–29^. To further explore the importance of cytoskeletal organization and activity in determining the nematic behavior of myogenic cells, we used sustained blebbistatin (5, 10, and 20 µM) treatments, an inhibitor of myosin-II, to perturb the actomyosin contractility and the organization of myoblasts and myotubes during the culture period (Extended Data Fig. 2, Supplementary Video 3). The length of blebbistatin-treated myofibers was significantly shorter than that of controls without having any significant change in cell density or fusion competent populations (Extended Data Fig. 2b-d). Importantly, despite myotubes still being able to form, the self-organization of the cell population was profoundly impaired in a dose-dependent manner, as illustrated by the significantly lower scalar order parameters (*S*) and higher nematic defect densities, compared to controls (Fig. 1g-h).

Blebbistatin-treated myoblasts displayed a higher density of topological defects (Fig. 1h), indicating that in this system, cytoskeletal maturation and organization dominate over active stress in determining the scalar order parameter (S) and defect density (Fig. 1g-h). This occurs through their modulation of orientational elasticity. Inhibiting myosin II with blebbistatin (5, 10, and 20 µM) produces two opposing mechanical effects. Reduced active stress generation would normally lower defect density^30^, whereas impaired cytoskeletal maturation decreases orientational elasticity, which favors defect formation^31^. The observed increase in defect density under blebbistatin treatment therefore suggests that reduced cytoskeletal organization has the stronger influence, underscoring the central role of cytoskeletal structure and activity in shaping the nematic properties of myogenic cells.

To separate the role of population wide actomyosin contractility from that of fusion-dependent myotube elongation and maturation, we inhibited fusion with EGTA (ethylene glycol-bis(β-aminoethyl ether)-N,N,N′,N′-tetra acetic acid), which blocks calcium-dependent membrane fusion between myocytes and myotubes without directly affecting contractility or differentiation^32^. Therefore, EGTA treatment markedly reduced fusion and resulted in shorter myotubes (Extended Data Fig. 2a-b), while preserving proliferation and differentiation (Extended Data Fig.2). Consequently, the order parameter evolved to values lower than control and comparable to those seen with 5 µM blebbistatin (Fig. 1g). This can be explained by the loss of long nematogens *i.e.,* long myotubes, which directly influence the orientational elasticity of the system. These results therefore indicate that actomyosin-driven contractility and fusion-mediated myotube growth together contribute to the maintenance and reinforcement of nematic self-organization during myogenic maturation with the evolution of actomyosin contractility being the main factor controlling the nematic behavior of the system.

Overall, these data show that populations of primary muscle cells exhibit characteristics of an active contractile nematic. Importantly, the system evolves/matures over time, leading to higher ordering while complex cellular events take place, such as proliferation, migration, fusion, and the massive growth of myotubes. In addition, such behaviors appear highly dependent on the cytoskeletal activity of these cells. Given that active nematic behavior is known to influence various biological processes, it prompted us to consider its potential implication in myoblast fusion.

### Fusion localization is affected by stable topological defects

To explore the temporal evolution of nematic organization and its potential link with fusion, we used chicken myoblasts transduced with lentiviruses encoding F-actin and non-muscle Myosin-II light chain (Myl9), each tagged with a fluorescent protein and driven by a strong ubiquitous promoter^33^. Additionally, low doses of Hoechst were added to the culture medium to allow for live staining of cell nuclei over 48h (Fig. 2a, Supplementary Video 4). The advantage of labeling the cytoskeleton with fluorescent reporters was two-fold. First, it allowed an easier detection of the fusion events. Indeed, whereas the transfer and alignment of new nuclei into nascent myotubes may be difficult to detect in very dense cell cultures, we could pinpoint fusion events unequivocally by detecting the mixing of the fluorescent signal between invading myoblasts and receiving myotubes, indicating the merging of their cytoskeletal proteins. (Fig. 2b, Extended Data Fig. 3a-b, Supplementary Video 5). Noteworthy, the fusion was often preceded by the formation of actin-rich foci resembling the F-actin-enriched invasive structures with comparable foci size and lifetime to the ones that were shown to mark the site of myoblast fusion in *Drosophila* and zebrafish^34,35^ (Fig. 2b, Extended Data Fig. 3a-d, Supplementary Video 6). Second, labeling the cytoskeleton with fluorescent reporters also allowed easy detection of the orientation field necessary to localize topological defects during the time course of the culture (Fig. 2c, Extended Data Fig. 4a-b).

**Fig. 2:**
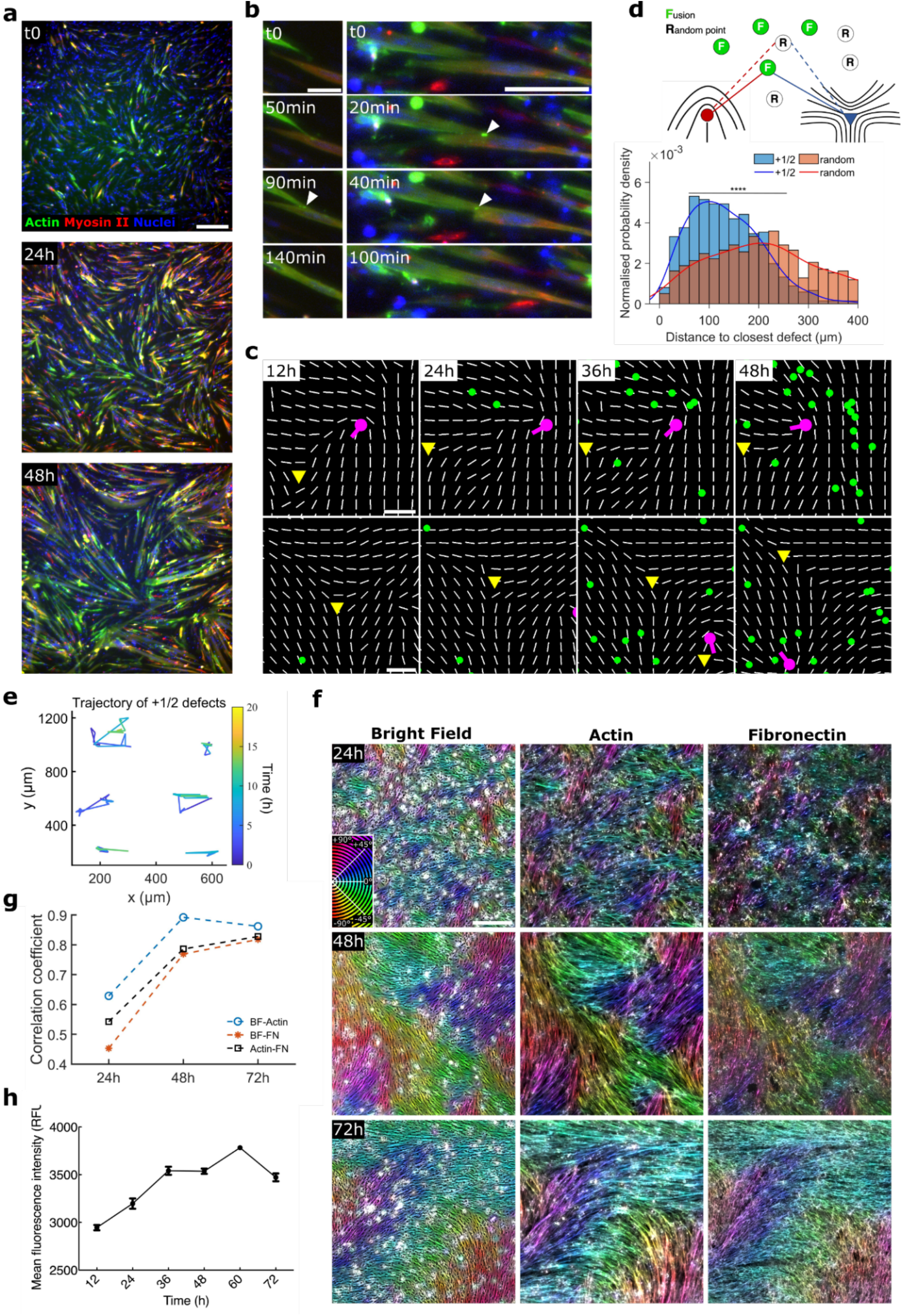
Topological defects affect the localization of myoblast fusion. **a**, Snapshots of the evolution of the myoblast culture transduced with the lentiviral construct allowing to observe actin (green) and myosin-II (red) cytoskeleton along a 48h period. Nuclei are stained with Hoechst (blue). Note the transition from myoblasts to myotubes. Scale bar, 200 µm. **b**, Close-ups on areas showing fusion events during the time-lapse. White arrowheads indicate the site of fusion characterized by an actin focus preceding cytoskeletal merging and fluorescence mixing. Scale bar, 50 µm. **c**, Time projection of the coordinates of the fusion events (green dots) that took place in proximity to a +1/2 defect (top row, magenta) or -1/2 defect (bottom row, yellow triangle) during a 48h movie. White lines illustrate the average orientation of the cells. Scale bar, 100 µm. **d**, Top: Schematics illustrating the measurement of the distance between fusion or random points and either to their closest +1/2 or -1/2 defect. Bottom: Probability density vs distance to closest defect and on top kernel density (continuous lines) to observe fusion events around +1/2 topological defects in comparison to random points (n = 678 fusion events and random points, 8327 +1/2 defects over 3 independent movies). A significance test was done on the raw data with two sample Kolmogorov–Smirnov test (p < 10⁻^4^). **e**, Trajectory of several +1/2 stable defects. **f**, From left to right, Bright-field, phalloidin, and fibronectin channel of primary myoblasts cultures fixed after one, two or three days in culture. Images are pseudo-colored according to local cell orientation. Scale bar, 100 µm. **g**, Orientational alignment (correlation coefficient) between bright-field (BF), actin, and fibronectin (FN) channels at various time points in culture. **h,** Mean fluorescence intensity of the fibronectin staining taken at various time in culture and measured with Fiji (n=3, 5 by 5 mosaics at 10x magnification from independent experiments for each time point, error bars are S.E. from the mean).

By comparing the positions of detected fusion events with the nematic organization of cells, we discovered that the probability of fusion events was highest towards the head of +1/2 defects (Fig. 2c-d, Extended Data Fig. 4b-c). In contrast, the probability of fusion events was lowest near -1/2 defects, which appeared even less favorable to fusion than the ones observed with randomly generated coordinates similar to “fusions” (Fig. 2c-d, Extended Data Fig. 4b-c). A majority of the fusion events took place during the second day of culture, at a time when more than half of the defects have already disappeared (Extended Data Fig. 4d-e). Strikingly, a subset of defects appeared very stable in space and time, with a lifetime that extends throughout the entire duration of the experiment (Fig. 2c, 2e). Moreover, we found a strong fusion accumulation around stable +1/2 defects over time (Extended Data Fig. 4b-c). As maturation of the system occurred, and elongated myotubes emerged, the size of the stable defects progressively increased (Fig. 1c). The stabilization of topological defects over time contrasts with the theoretical maturation of active nematic systems, which exhibit ’active turbulence’, a chaotic state characterized by the constant generation and annihilation of topological defects^23^. To explain the emergence of stable defects in our system, we hypothesized that the stabilization mechanism can arise from changes at the cell–substrate interface, particularly through extracellular matrix (ECM) remodeling^36–38^. Consistent with this idea, fibronectin immunostaining revealed progressive, cell-driven ECM deposition and restructuring, accompanied by the development of an ECM orientation field that gradually evolved to closely match the cellular orientation inferred from brightfield and actin imaging (Fig. 2f-h). Such ECM remodeling could substantially slow or constrain defect motion, thereby promoting their stabilization. In addition, the patterned ECM architecture generated by the cells may guide their migration along defect lines (Extended Data Fig. 5a-c). This could potentially enhance the likelihood of fusion events at specific locations through cell stabilization. Therefore, we quantified cell speed as a proxy for contact duration and we observed comparable cell speed in proximity to each type of defect’s core or far away from them (Extended Data Fig. 5d). Moreover, directly following the duration of a subset of cell-cell interactions showed no significative difference between +1/2 and -1/2 defects (Extended Data Fig. 5e). Thus, the increased fusion frequency at +1/2 defects cannot be accounted for by longer or more stable cell–cell contacts and is therefore the consequence of other mechanisms.

Altogether, these observations provide compelling evidence that the presence of topological defects significantly influences the localization of fusion events, defining stable +1/2 defects as central hubs where fusion preferentially takes place.

### Fusion at +1/2 defects correlates with actomyosin-driven compressive stresses

It has been long known that the rate of fusion is proportional to the density of myoblasts *in vitro*^39,40^. It was thus possible that the significant increase in fusion events associated with +1/2 topological defects would be due to a higher density of cells in these regions. To determine whether this was the case, we quantified cell density in the vicinity of fusion events or around topological defects over time, and normalized this against the cell density in the entire field of view (Fig. 3a, see methods section). These analyses were performed in the first half of the experiments, which covers the earliest fusion events, thereby avoiding a possible bias due to the emergence of massive myotubes. We did not observe any significant change in cell density in either of the cases, suggesting that, within our experimental setting (where cells are at confluency soon after the start of the experiment), fusion events do not depend on local cell density enrichment but rather on cell organization. Similar results for cell density were obtained when analyzing -1/2 defects in the first half of the experiments (Extended Data Fig. 6a), or both defect types over the entire duration of the movies (Extended Data Fig. 6b-c).

**Fig. 3:**
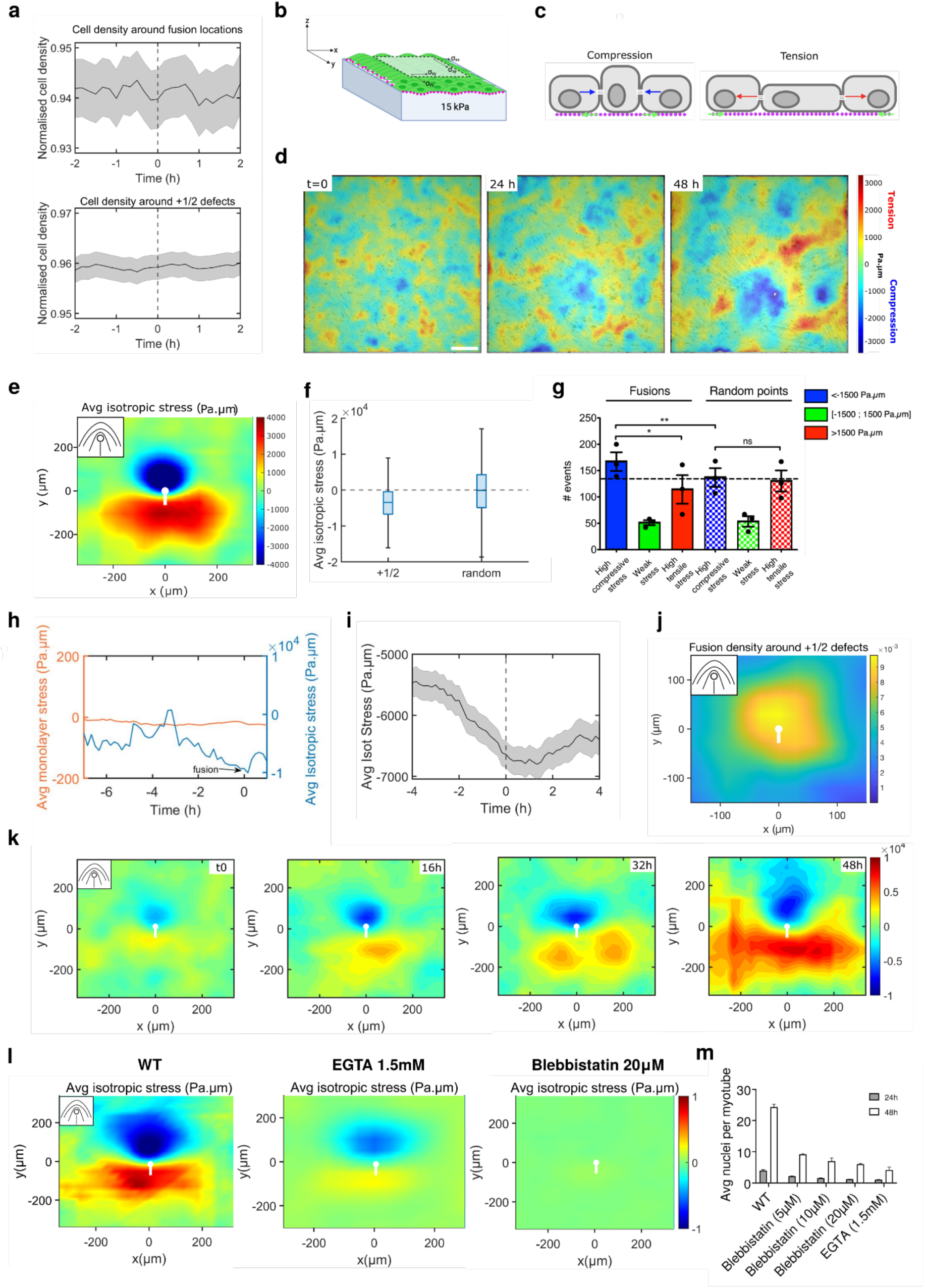
Fusion is enriched in areas of compressive stress progressively generated at the head of +1/2 defects. **a**, Averaged evolution of the cell density around fusion events (top) and the core of +1/2 defects (bottom). t0 corresponds to the time of the fusion event or the birth of the defect respectively (analysis was performed over the first 24h of 3 independent 48h movies, error bars are S.E. from the mean). **b**, Schematic of BISM utilizing data from traction force microscopy to infer the isotropic stress in cell monolayers (details in methods). **c**, Schematic illustrating the tension and compression experienced by the cells as a result of their activity and neighbor interaction (blue and red arrow), and the corresponding displacement of the beads attached to the substrate (green arrows). **d**, Representative stress map obtained for a full movie overlaying the bright-field image of the cells. Yellow to red regions (positive values) indicate areas of tension while areas of compression are represented in blue (negative values). Scale bar, 200 µm. **e**, Averaged isotropic stress map around +1/2 defects (n = 8327 +1/2 defects in 3 independent movies). **f**, Stress intensities around +1/2 defects and at random locations (n=998). **g**, Bar plot illustrating the distribution of the fusion events or random points according to the averaged stress in 50×50 pixels regions (32,5×32,5 µm) around each event (n = 987 fusion events and random points from 3 independent movies, error bars are S.E. from the mean). P values were calculated using paired two-tailed t-test *P<0,05 and **P<0,01. **h**, Representative time evolution of the average isotropic stress around a fusion event showing occurrence of the fusion at high compressive stress (blue) compared to average isotropic monolayer stress (yellow). **i**, Evolution of Average isotropic stress around fusion locations (50×50 pixels regions) occurring in compressing regions, t=0 indicates the occurrence of the fusion event. **j**, Density map of fusion events around +1/2 defects (incorporate every fusion event falling in the selected area around the core of the defects over 3 independent movies, n = 719 fusion events and n= 8327 +1/2 defects). **k**, Evolution of average isotropic stress maps obtained for +1/2 defects during the culture period (averaged at each hour over 3 independent movies). **l,** Average isotropic map obtained for +1/2 defects for WT, EGTA (1.5mM), Blebbistatin (20µM) (normalized with respect to control) (averaged for 3 independent movies), **m**, Progression of myotube lengths with various drug treatments at 24h and 48h time points (for 3 independent movies, error bars are S.E. from the mean). **e**, **j**, **k, l** Inset is a schematic of the orientation field characteristic of +1/2 defects and central mark represents the core of the defect.

Since local density was not the driver of fusion events, we reasoned that mechanical stress could play a major role. Topological defects have already been described as hotspots of mechanical stresses^13,14,41^. We characterized the stress patterns in our experimental setting using Bayesian inversion stress microscopy (BISM)^42,43^ (Fig. 3b-3c). We first considered the spatial distribution of the isotropic stress, and observed heterogeneous patterns including regions of tensile (positive) and compressive (negative) mechanical stress (Fig. 3d). We then examined the stress patterns associated with the topological defects, and found that +1/2 defects displayed compressive stress in their head region, in close proximity to the defect core, and tensile stress in their tail (Fig. 3e, Extended Data Fig. 7a-b). When comparing global stress levels to those near defects, we found that the heads of +1/2 topological defects consistently corresponded to regions of maximal compressive stress (Fig. 3f, Extended Data Fig. 7c-d). By contrast, –1/2 defects displayed markedly weaker stress intensities, characterized by a tensile core surrounded by alternating tensile and compressive regions arranged in a three-foil symmetric pattern (Extended Data Fig. 7a-c, 7p). The high tensile stresses at the cores of –1/2 defects have been reported to induce cell escape^44^ or even tissue fracture^14^, providing a mechanistic explanation for why fusion is far less likely to occur near –1/2 than +1/2 defects (Fig. 2d, 3e-f, Extended Data Fig. 7a-d).

We therefore focused on +1/2 topological defects and asked to what extent the localized compressive stresses at their heads correlate with myoblast fusion. We found a significant enrichment of fusion events within regions experiencing high compressive stress (Fig. 3g–i). Density maps obtained by averaging fusion-event positions around +1/2 defects further showed that fusion is strongly concentrated in the compressive zone at the defect head (Fig. 3j, Extended Data Fig. 7b). Moreover, we observed that the stable +1/2 defects showing the highest compressive stress are also the ones concentrating the most fusion events (Extended data Fig. 7e).

To further probe the temporal relationship between mechanics and cell fusion, we performed a temporal correlation analysis between local stress and fusion occurrence around stable defects. We first observed a substantial negative correlation between compressive stress and the timing of fusion events (Extended Data Fig. 7f-g). Then, time-resolved analyses revealed that increases in compressive stress systematically precede fusion: the distribution of time differences between peak stress and subsequent fusion events shows a pronounced peak at negative time delays (Extended Data Fig.7h-l). Notably, similar compressive stress patterns around +1/2 topological defects have been reported in other active nematic systems, including MDCK, MCF-7, and E-cadherin–deficient epithelial cells^13,26^, as well as in monolayers cultured on soft substrates^14^, where no fusion occurs. This supports the conclusion that the observed stress configuration is an intrinsic property of active nematic organization rather than a consequence of the fusion process itself. One could also ask whether differentiation patterns could underlie the spatial bias in fusion. We therefore quantified Myogenin (MyoG) expression, which marks fusion-competent cells upstream of fusogen expression, and found that MyoG-positive cells were distributed homogeneously across the culture at multiple time points (Supplementary Fig. 1). These data argue against spatial bias in differentiation or fusogen expression as the driver of the stress–fusion correlation. Altogether, these observations establish that active nematic organization generates the compressive stresses associated with comet-shaped defects, and that theses stresses generated upstream of fusions events appear to strongly bias the spatial localization of myoblast fusion independently from their differentiation process.

Remarkably, close observation of the evolution of overall isotropic stress landscape revealed a major increase in magnitude in both tensile and compressive regions over time, with values exceeding ± 7000 Pa.µm (respectively), as well as an enlargement of their domain sizes (∼ 4-fold, Extended Data Fig. 7m-o, Supplementary Fig. 2). Most importantly, we observed a strong evolution of the defect stress patterns. Even though the overall compressive-tensile stress pattern remains, we observed a significant increase in stress intensity and area covered by the pattern over time for both defect types (Fig. 3k, Extended Data Fig. 7p, Supplementary Video 7-8). The stress evolution pattern correlates with the maturation process, as myotubes continue to fuse and grow around defects (Extended Data Fig. 7o). Together, these findings establish that myogenic differentiation is accompanied by a dynamic strengthening of defect-associated stresses, revealing a fundamental remodeling of tissue mechanics as myotubes emerge.

In order to explore the relative participation of cell contractility, as well as fusion-driven nematogen growth (i.e., myotube growth) in the evolution of the nematic and stress landscape, we performed experiments on cell-generated stresses around topological defects under conditions of contractility inhibition—using blebbistatin (5, 10, 20 µM)—and fusion inhibition through calcium chelation with EGTA (1.5 mM). Increasing blebbistatin concentrations progressively reduced mechanical forces, and at 20 µM, contractility was effectively at minimum level, therefore highlighting again the key role of the cytoskeletal activity in the evolution of the system. At the highest dose of blebbistatin, the average number of nuclei per myotube, a proxy for cumulative fusion, dropped to roughly 25% of control levels (Fig. 3l–m, Extended Data Fig. 8). These results indicate that while a minimal level of fusion can still occur, the vast majority of fusion events were abolished; probably due to a combination of effects of the treatment on actomyosin contractility at the cellular scale, where it affects stress generation and nematic behavior, and at the subcellular scale where it can alter the key actin-dependent steps of the fusion machinery. The role of actomyosin in stress generation becomes even more evident when contrasted with EGTA-induced fusion inhibition: although EGTA leaves actomyosin activity intact, it eliminates elongated myotube formation despite preserving stress levels at roughly 65% of control (Extended Data Fig. 8). Notably, these stresses remain substantially higher than those observed at any blebbistatin dose (Fig. 3l–m, Extended Data Fig. 8). The higher stress levels observed in EGTA-treated cultures compared with blebbistatin-treated cultures can be explained by the preservation of contractility in most of the cell population. By contrast, the approximately 35% reduction in stress intensity relative to control conditions likely results from the impaired formation of long myotubes due to blocked fusion. Under EGTA treatment, the myogenic population therefore lacks long, contractile myotubes, whose mechanical contribution cannot be compensated for by mononucleated cells or very short myotubes. Together, these comparisons support the conclusion that actomyosin contractility is the main driver shaping the nematic behavior of the cells.

### Stable topological defects amplify the extent and intensity of compressive regions

The fusion process leads to the formation of very long myotubes, sustained by a scaffold of longitudinal actin bundles spanning the length of myotubes, the activity of which is essential for maintaining the shape and physical properties of the myotubes, as shown by blebbistatin treatment^27–29^ (Extended Data Fig. 2). Thus, the dramatic growth of myotubes is likely to significantly impact the physical properties of the tissue, such as orientational elasticity, which in nematics is expected to increase with the increasing length of the nematogens^45^. To better understand the relationship between defects, fusion and maturation process, we hypothesized that stabilization of mechanical stresses around topological defects was a crucial parameter. We thus turned to a continuum model of active nematics^46^ with monotonically increasing orientational elasticity to mimic evolution of the cytoskeleton during maturation and growth of myotubes, culminating in a high-elasticity regime. Before incorporating the effects of fusion and ECM coupling, we first tested whether the intrinsic active nematic behavior alone could account for the observed stress distribution. By explicitly defining bend and splay components of orientational elasticity (Extended Data Fig. 9a), the model reproduced the same compressive and tensile stress patterns around topological defects even in the absence of fusion, indicating that these stresses emerge as a fundamental property of the active nematic organization rather than as a byproduct of fusion events. The synergistic effects of activity and time-dependent elasticity were sufficient to qualitatively capture the drop in defect density (Extended Data Fig. 9b-d) and the stress patterns around topological defects (Fig. 4e). However, we still had to explain the stabilization of topological defects and the significant amplification in size and intensity of stresses around defects (Fig. 3e, 3k, Extended Data Fig. 7p). Thus, we added another level of complexity to this minimal model, taking into account the experimental observation that ECM patterning closely followed the orientation of primary myoblasts over time (Fig. 2f-g). We therefore hypothesized that the cell-induced, dynamically evolving ECM plays a major role in stabilizing defect motion over time through creating an increasing friction to the motion of cells. To account for this experimental observation, we thus introduced a temporally evolving orientation field for the ECM, which followed the evolution of the cell orientation field with a time delay, and resulted in a time-dependent friction experienced by cellular active nematics (Fig. 4a-b, see methods for details of the model). We observed that as the simulation time progressed, drastic changes in the features of the system emerged (Supplementary Video 9). First, the number of topological defects (Extended Data Fig. 9c) decreased, in agreement with experimental data (Extended Data Fig. 9d). This was accompanied by a sharp reduction in defect velocity (Fig. 4c), which corresponded to the stabilization of defects in the experimental data (Fig. 2c, 2e, 4d). The simulation also reproduced the flow (Extended Data Fig. 9e), the strain rate patterns around the defects (Extended Data Fig. 9f), as well as the decrease in defect density (Fig. 4a). More importantly, calculating the average stress around defects revealed that the compressive region around the head of the +1/2 defects increased significantly in both area and intensity (Fig. 4e-g). This significant amplification of the extent and intensity of the compression around defects with time was only observed when the coupling to the dynamically evolving ECM is accounted for, highlighting the importance of the active crosstalk between the cell layer and the ECM in shaping the collective cell organization and the mechanical stress intensity patterns.

**Fig. 4:**
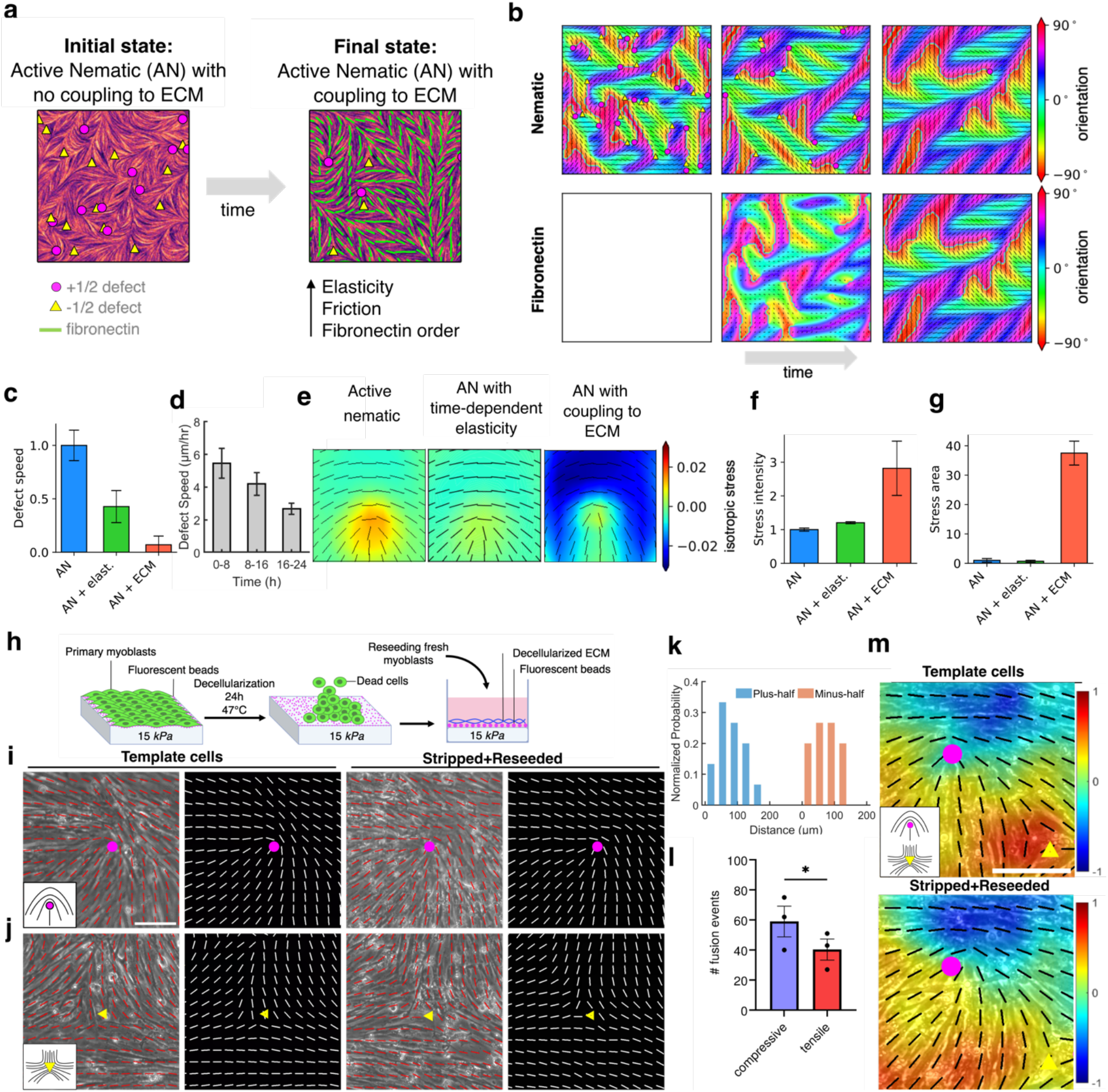
Change in the orientational elasticity and coupling to ECM dictates the stress patterns around defects. **a**, Schematic representation of the modelling framework with time-dependent orientational elasticity and ECM evolution. Once system reaches active turbulence at t0, the elasticity, friction and fibronectin order increase monotonically until saturation at maximal value. **b**, Simulation snapshots of nematic (top) and fibronectin (bottom) orientation at different times of the simulation. **c**, Average +1/2 defect speed for active nematic (AN) with no coupling to ECM, AN with time-dependent elasticity, AN with time-dependent elasticity coupling to ECM. **d**, defect speed from experiments at various time periods during the culture period (n=10 stable defects). **e**, Average isotropic stress around +1/2 defect in active nematic (AN), AN with time-dependent elasticity and AN with time-dependent elasticity coupling to ECM. **f–g**, Stress intensity (**f**) and area (**g**) at the head of the +1/2 defect in active nematic (AN), AN with time-dependent elasticity and AN with time-dependent elasticity coupling to ECM. **h,** Schematic of stripping/reseeding procedure: After nematic organization and traction fields were established, myoblasts were removed while preserving the underlying ECM, and fresh myoblasts were reseeded onto the stripped matrix. **i-j,** Snapshots of cultures before stripping (Template) and after reseeding (Stripped+Reseeded) showing the reappearance of plus half (**i**) and minus half (**j**) nematic defects at the same locations as in the template culture. Alternating panels show the corresponding orientation vector fields. **k,** Normalized probability density of fusion events as a function of position around +1/2 and –1/2 defects after reseeding. **l,** Number of fusion events in compressive versus tensile regions after reseeding (averaged over 3 independent movies, significance test was set at p<0.05.). **m,** Isotropic stress map around defects in template (top) and stripped+reseeded matrix (bottom), overlaid with the averaged local orientation of the cells (black lines). **i**, **j**, **m**, magenta dot and yellow triangle respectively marks the core of +1/2 and minus half defects and insets are schematics of the orientation field characteristic of +1/2 and -1/2 defects. **c, d, f, g, l** Error bars represent standard error (S.E.) from the mean.

To validate the predictions of the model and strengthen our observation that fusion is enriched in regions of high compressive stress organized by active nematic behavior, we performed a decellularization assay in which cultured myoblasts were removed and new ones were seeded onto the ECM left behind by the original cells^47^ (Fig. 4h, Methods). After traction fields and nematic defects had formed, all template cells were stripped away while preserving the cell-generated matrix (Supplementary Fig. 3). Remarkably, the reseeded cells recapitulated the ECM-imprinted orientation field, and both +1/2 and -1/2 defects re-emerged at the same spatial locations (Fig. 4i-k). Fusion then occurred preferentially at these defect-associated stress sites, with a predictable bias toward regions under compression (Fig. 4l, 4m). These results demonstrate that the ECM alone retains the spatial template that governs nematic organization, stress patterning, and the associated spatially biased distribution of fusion events. Furthermore, as we showed previously that modulating cell contractility affects tissue-stress patterns, we decided to assess the state of the ECM following blebbistatin treatment. With sustained inhibition of contractility, the ECM becomes progressively disorganized in a dose-dependent fashion (Extended Data Fig. 10). Therefore, this illustrates that a minimal level of actomyosin activity is necessary for ECM patterning. This requirement may arise from a combination of direct traction forces exerted on the ECM and the maintenance of the elongated cell shape characteristic of myoblasts and myotubes. In addition, EGTA treatment led to a dramatic reduction in fusion, reflected in both the average number of nuclei per myotube and in myotube elongation (Fig. 3m, Extended Data 2b). By contrast, the cells’ ability to self-organize and to remodel the ECM over the course of the culture period was only mildly affected (Fig. 1g, Extended Data Fig. 10). EGTA also caused a smaller decrease in stress intensity than any blebbistatin treatment (Fig 3l, Extended Data 8). These results suggest that the observed delay in the orientational order and stress evolution arises primarily from the loss of long, fused myotubes and of their mechanical activity, supporting the involvement of orientational elasticity in the self-organization process modeled computationally. Moreover, they reinforce the conclusion that ECM deposition and shaping driven by actomyosin contractility are central to generating and organizing mechanical stress. Together, these observations support the conclusion that ECM patterning and actomyosin contractility jointly contribute to the emergence of stress fields and to the organization of myotube formation. Collectively, these complementary experiments move beyond simple correlation and support a model in which ECM remodeling is necessary and sufficient to stabilize defect positions, the associated mechanical stresses and generate the observed bias in fusion locations.

Together, our modeling and experimental data allow us to decipher the main nested mechanisms ruling the evolution of this system, revealing a unified mechanism by which time-dependent changes in myoblast physical properties orchestrate stress patterning and fusion. The continuum model shows that the interplay between cellular activity, increasing orientational elasticity during myotube maturation, and rising friction from a dynamically evolving ECM is sufficient to spatiotemporally organize stresses around topological defects. This coupling generates persistent compression hotspots whose size and intensity grow over time. Consistent with these predictions, our experiments identify stable defect-associated compressive zones, an ECM architecture that mirrors and reinforces the local nematic organization, and a marked enrichment of fusion events at these mechanically stressed sites. Together, these findings demonstrate that ECM remodeling is both necessary and sufficient to stabilize defect positions and that the stresses concentrated around these defects appear to function as mechanical organizers of fusion and collective tissue patterning.

## Discussion

In the present study, the active contractile nematic behavior exhibited by primary embryonic chicken myoblasts in culture directs patterns of self-organization and apparently promotes fusion in regions of high compressive stresses. Interestingly, these regions correspond to the localization of comet-shaped (+1/2) topological defects. This enrichment was particularly pronounced in the head region of these defects, where it correlated with elevated compressive stress^13,14,41^. On the other hand, -1/2 defects, where tensile stresses are largely dominant, were found to be less favorable for fusion.

As earlier studies^39,40^ demonstrated that cell density is an important factor determining the rate of fusion *in vitro*, it was hypothesized that density might affect the initiation of fusion by increasing the encounter of cells competent to fuse. Even though we observed, as most researchers working with myoblasts did, that cell confluency is essential to fusion initiation, our data demonstrate that when fusion is triggered, fusion hotspots are independent of cell density enrichment. In contrast to previous studies^16^, despite the cells being at confluence, we did not observe notable change of density around early fusion events nor around the core of +1/2 defects where fusion is enriched. This suggests that cellular density alone does not account for fusion probability in our system, or for the progressive increase in stress we observed. Instead, our findings highlight the major role of mechanical stresses, particularly the ones around +1/2 topological defects, as organizers favoring myoblast fusion.

Previous studies^13–16,37,44^ have highlighted the critical role of active nematics and topological defects in regulating biological processes both *in vitro* and *in vivo*. Building on this foundation, our work uniquely examines the dynamic evolution of the system’s intrinsic properties. As myoblasts fuse to form elongated myotubes, we observe an enhancement in nematic ordering alongside a reduction in topological defects. By contrast, inhibiting cellular activity through myosin II suppression leads to an increase in topological defects. Additionally, we discovered that the spatial organization of cells is mirrored by that of the extracellular matrix (ECM) over time and that myosin II inhibition triggered ECM disorganization. Our results indicate that actomyosin contractility plays a central role in sustaining nematic self-organization in myogenic populations. Rather than acting as an isolated determinant, contractility contributes through a combination of effects on cell elongation and remodeling of the extracellular matrix (ECM), which together maintain the mechanical coupling required for the emergence and persistence of ordered patterns. These findings highlight how contractility-dependent mechanics, acting at the supracellular level, shape the collective behavior of myoblasts. Furthermore, inhibition of fusion-driven myotube growth through calcium chelation led to a delay in the ordering process without stopping it totally and a notable decrease in mechanical stresses. These findings highlight the critical role of dynamically evolving orientational elasticity, emerging from actomyosin network maturation and myotube growth, in driving cell self-organization during myoblast differentiation and myotube formation.

Furthermore, mathematical modeling demonstrates that the interplay between cellular dynamics and ECM reorganization is central to understand the stabilization of topological defects observed in our study, as well as the stress amplification mechanism that apparently promotes fusion events in the growing compressive stress regions associated to +1/2 topological defects. The importance of time-dependent physical changes in active materials, along with the positive feedback between passive (ECM) and active (cells) systems^48^, highlights a fundamental mechanism that could reshape our understanding of tissue formation and regenerative processes.

Integrating modeling, experimental observations, and comparative analyses, we identify a two-component model in which the canonical fusion machinery still sets the baseline fusogenic potential, while cell activity and ECM-related stresses apparently potentiate and spatially pattern fusion across the tissue. Consequently, stress-dependent fusion could emerge as the principal driver of fusion organization.

Our experiments provide functional evidence that strengthens the mechanistic link between nematic organization, ECM alignment, compressive stresses, and fusion localization. Together, they position our mechanical model firmly within the established framework of muscle biology and open a new research avenue in which the supracellular stress landscape contributes to efficient, spatially patterned myoblast fusion. *In vivo*, however, it remains unclear whether topological defects emerge prior to myotube ordering. Embryonic muscle develops within a mechanically heterogeneous environment shaped by neighboring tissues, spatially patterned ECM, and dynamic forces. Thus, even in the absence of clearly identifiable defects, tissue-scale mechanical stresses are likely to contribute to the spatial regulation of fusion during development. Overall, these insights highlight the central role of tissue-scale mechanics in shaping the emergence and spatial patterning of muscle during development.

Our results emphasize the pivotal role of active nematics and mechanical stresses in unraveling the intricacies of complex processes of morphogenesis, such as the self-organization and development of primary muscle progenitors. They show the critical importance of dynamically evolving material properties during myoblast to myotube transition. This evolving framework underscores the dynamic nature of biological systems, where unique mechanical properties can trigger cascading biological events, especially during processes of morphogenesis^49^. Our study brings new insight in the biophysical regulation of myoblast fusion and raise intriguing questions regarding the contribution of mechanical stresses, especially compressive stress, to the spatiotemporal regulation of myogenesis. These findings pave the way for innovative approaches to study and potentially modulate complex morphogenic events in various biological contexts such as embryonic development, tissue regeneration or organoid development.

## Materials and Methods

### Chicken primary embryonic myoblasts culture

Skeletal muscle progenitors were isolated from 11-days-old WT chicken embryos. Briefly, the hindlimbs muscles were removed and minced to pulp in a sterile Petri dish using dissection scissors. The pulp was then incubated 12 minutes at 37°C in Trypsin-EDTA (Thermo Fisher) diluted to 0.125% in PBS accompanied by two trituration steps. The trypsin was then inhibited by the addition of fetal bovine serum (FBS) and discarded after cell centrifugation. The cell pellet was resuspended in growth medium composed of Dulbecco’s Modified Eagle Medium (DMEM) GlutaMAX^TM^ (Thermo Fisher) supplemented with 20% FBS and 1% Penicillin/Streptomycin (P/S; Thermo Fisher), and filtered through a 70 μm nylon cell strainer to allow the isolation of mononucleated cells from muscle fibers and debris. The cell suspension then underwent two one-hour rounds of pre-plating on uncoated Petri dishes to deplete the suspension from fibroblasts and red blood cells attaching quickly to the plate. Un-attached cells were then collected, centrifugated and resuspended in growth medium before plating at 70% confluency for each experiment. Cells were seeded on fibronectin (15 μg/ml; Merck) coated soft silicone gel PDMS (polydimethylsiloxane; Dow Corning) substrate with a stiffness of 15 *k*Pa in 24 wells plates.

### Lentiviral production

Lentiviral particles containing the CAGGS-LifeactNeonGreen-IRES-tdTomatoMyl9 vector construct were produced in HEK 293T cells. The cells were plated in 75 cm^2^ cell culture Petri dishes in DMEM GlutaMAX^TM^ complemented with 10% FBS and 1% P/S for one day. The medium was then switched to DMEM GlutaMAX^TM^ with 5% FBS and transfected with 1,5 ml of OptiMEM (Thermo Fisher) with 0,04% Polyethyleneimine (Merck) containing 6,65 µg of vector, 4,2 µg of psPax2 and 2,7 µg of pMD2.G plasmids overnight at 37°C and 5% CO2. psPAX2 and pMD2.G were gifts from Didier Trono (Addgene plasmid # 12260; http://n2t.net/addgene:12260; RRID:Addgene_12260; Addgene plasmid # 12259; http://n2t.net/addgene:12259; RRID:Addgene_12259). The next day, the medium was replaced with fresh DMEM GlutaMAX^TM^, 5% FBS, 1% P/S for 24h, after which the supernatant was collected and conserved at 4°C. This step was repeated a second time. Afterward, the collected medium was filtered using 0,22 µm non-pyrogeic syringe filters (Merck), and centrifugated at 1250 rpm over night at 4°C. The supernatant was then discarded and all the pellets resuspended in a small amount of DMEM GlutaMAX^TM^ and conserved at - 80°C until use.

### Lentiviral transduction

In order to observe the actin and Myosin-II cytoskeleton, primary myoblasts were transduced with lentiviral particles containing a vector coding for the expression of Lifeact-NeonGreen and tdTomato-Myl9 fusion proteins under the control of the ubiquitous CAGGS promoter (CMV/chick β-actin promoter/enhancer). Infection was initiated 6h after cell seeding for 24h in growth medium without antibiotics and with 5 μg/ml polybrene (Merck). The medium was then replaced for fresh growth medium with 1% P/S overnight to wait until sufficient expression of the lentiviral construct and then switched to differentiation medium composed of DMEM GlutaMAX^TM^ supplemented with 5% FBS and 1% Penicillin/Streptomycin, before live imaging.

### Cell treatments

In order to inhibit Myosin-II activity and observe the dose-response of cell activity, primary chicken myoblasts were switched to differentiation medium 6h after cell seeding and treated with 5, 10 or 20µM (-)-blebbistatin (Merck) or the equivalent volume of Dimethyl sulfoxyde (DMSO, Merck), as a control, before live imaging or fixation at 24h and 48h of culture.

In order to directly inhibit the fusion machinery, primary chicken myoblasts were cultured in a similar setting and treated with 1.5 mM of the calcium chelator EGTA (Merck) diluted in distilled water.

### Live cell imaging

Transduced cells were imaged during TFM experiments for 48h after switch of the culture to differentiation medium with addition of Hoechst 33342 (Thermo Fisher, Table1) at 0,02 μg/ml to stain the nuclei. The cells were recorded every 10 minutes under wide field microscope Axio Observer Z1 (Zeiss) equipped with a sCMOS Orca Flash4 LT camera (Hamamatsu) at 10x magnification (10 X Pl NeoFluar ON 0.5 Ph). For the duration of the time lapse, the cells were kept at 37°C and 5% CO2 using an incubator chamber (Life Imaging Service).

For phase contrast large field live imaging, primary chicken myoblasts where imaged for a duration of 48 to 72h every 20 minutes at 10X magnification (10 × 0.25LD Ph) to obtain 25-tile mosaics, using a Zeiss Axio Observer 7 equipped with an Orca R^2^ camera (Hamamatsu) and incubator chamber. Live imaging was initiated 6h after cell seeding. Culture medium was switched from growth to differentiation medium just before imaging. For high resolution imaging, transduced cells were recorded every 4 minutes during 24h under Nikon spinning disk (CSU W1) microscope with 40x water objective.

### Stripping and reseeding assay (decellularization)

Primary chicken myoblasts were cultured on 15 *k*Pa PDMS and recorded live for 48h after switch of the culture to differentiation medium. Following initial culture, the template cells culture was decellularized by adapting the heat shock treatment from^47^ to preserve the extracellular matrix derived from template cells. For this, the cells were kept in PBS and incubated for 24h at 47°C to induce cell death. Following this incubation, the decellularized ECM was meticulously washed with PBS before the seeding of fresh primary chicken myoblasts which were then cultured and recorded in a similar manner to the template cells. This protocol was also combined with TFM associated to transduction of both the template cells and the ones used for recellularization with the lentivirus allowing actin and myosin visualization as well as Hoechst nuclei staining. When performing these experiments, both batches of cells were strictly imaged in the same area of each well to ensure comparable organization and traction force measurements between cell batches. Moreover, when performing TFM in this setting, the reference image allowing traction force measurement was taken at the end of the recording session of the second batch of cells seeded on the decellularized ECM.

### Fusion events quantifications

Manual counting of fusion events was performed using Fiji multi-point tool^50^, based on identifying actin foci characteristic of fusion synapses and cytoskeletal signal mixing events indicative of cytoplasmic coalescence. The heterogenous expression of the lentiviral construct among cells was essential for pinpointing the precise timing of cytoplasmic coalescence by allowing visualization of the cytoskeletal signal dilution from one cell into its fusion partner.

### Immunofluorescence

Cells were fixed with 4% formaldehyde in PBS for 10 minutes and washed three times with PBS. Permeabilization and blocking was performed by incubating in washing buffer solution (2% BSA, 0,1% Triton X100, 0,0002% SDS) of 1 hour. Cells were then incubated overnight at 4°C with primary antibodies (myogenin monoclonal antibody, 1:1000 dilution, F5D-S, Developmental studies hybridoma bank (DSHB); myosin heavy chain 1E monoclonal antibody; 1:10 dilution, DSHB; fibronectin monoclonal antibody, 1:100 dilution, B3/D6, DSHB) diluted in washing buffer. After three 10-minutes washes in washing buffer, cells were incubated for 1 hour at room temperature with secondary antibodies and dyes (Phalloidin-488, 1:200 dilution, A12379, Thermo Fisher Scientific; DAPI, 1:10000 dilution, D1306, Thermo Fisher Scientific; Hoechst 33342, 1:500000 dilution, H3570, Thermo Fisher Scientific; mouse IgG2a-555 antibody, 1:1000 dilution, A21137, Thermo Fisher Scientific; mouse IgG2b-647, 1:1000 dilution, A21242, Thermo Fisher Scientific). Finally, cells were washed three times for 10 minutes with washing buffer and stored in PBS until imaging.

### Nuclei and myotubes segmentation and analysis

Automatic nuclei segmentation was performed using a custom model for the StarDist2D^51^ Fiji plugin. Model training was done on the ZeroCostDL4Mic google collab^52^ based on manually generated training and test pre-segmented samples to segment nuclei in live recordings. Myotube characteristics were measured on of 10x mosaic images following MyHC staining. Myotubes were assessed by manually segmenting each individual myotubes as a freehand line ROI in Fiji. Myotube length was derived from the ROI measurements. Average number of nuclei per myotube was obtained from the mask of segmented myotube and finding the number of nuclei falling inside the mask using a custom-built macro in Fiji.

### Fibronectin deposition analysis

To estimate fibronectin deposition, cells were fixed at various times, stained with anti-fibronectin antibody, and 25-tile mosaics were acquired using a 10x objective. Total fibronectin signal intensity was measured in Fiji.

### Automated nematic characterization

To obtain the nematic director field, we employed the strategies described elsewhere^13^. In short, we used a phase contrast image or actin-myosin fluorescence channels to obtain the nematic director field. The images were smoothed using Bandpass Filter in Fiji. Fiji plugin OrientationJ was then used to detect the direction of the largest eigenvector of the structure tensor of the image^53^ for each pixel. The local nematic order parameter tensor Q was calculated for each point on a grid that discretized the image, using an in-house Matlab code, and allowed the representation of the corresponding nematic directors on the images. Automatic nematic defect detection was done based on calculation of winding number^54^ and primarily two types of defects (+1/2 and −1/2) were observed. The identified defects present for at least 0.5-1hr at the same location were considered for analysis. Density of nematic defects was calculated by dividing their average number at each time point by the area of the field of view. To determine a measure for the global ordering, orientation field was then divided into local regions containing about 16-20 nematic directors. The global ordering of the cell population was then measured by averaging the local regions to obtain the scalar order parameter (*S*) that represents the alignment of the nematic directors in all local regions.

### Defect and cell trajectory tracking

The trajectory of individual defects and cells was manually tracked using the Fiji plugin TrackMate^55^.

### Correlation coefficient analysis (BF-Actin-Fibronectin)

The orientation of three channels (BF, Actin, Fibronectin) was obtained using OrientaionJ in Fiji. To obtain the correlation coefficient, the correlation between obtained orientation angles were performed on the whole image for all the time points using MATLAB.

### Traction Force Microscopy (TFM)

Myoblasts were seeded on soft PDMS substrate with attached fluorescent beads (200 nm, Invitrogen). To prepare the PDMS with a stiffness of 15 *k*Pa, CyA and CyB components (Dow Corning) were mixed at 1:1 ratio and manually spin-coated on 24 wells plates to achieve a flat substrate. After that, the substrate was cured for 2h at 80 °C and then was silanized with 10% 3-aminopropyl trimethoxysilane (Sigma) in ethanol for 5-10 min. Carboxylated fluorescent beads (200 nm, Invitrogen) were functionalized on the substrate at 1:500 dilution in deionized water. The substrate was coated with fibronectin (15 μg/ml) and incubated on the substrate for 2 h prior to myoblasts seeding. To obtain the resting position of the beads, the cells were completely removed by adding 10% SDS. Beads displacements (with respect to its resting state), acquired during experiments, were measured and converted to cell traction forces^56^.

To obtain the traction forces, the reference beads image was first concatenated with the beads images containing cells, such that the reference frame preceded the beads image with cells using home developed Fiji macro. The images were then stabilized using the Image stabilizer^57^ or template matching^58^ plugin within Fiji to correct for any x-y drift, following which, intensity correction was applied to these images if needed. The velocities of beads movements were obtained using PIV (Particle Image Velocimetry) in MATLAB^59^. Using the FTTC^60^ plugin, we obtained traction force fields from beads velocities using a regularization parameter of 9 × 10^−9^. From the traction force fields, we estimated the stress using BISM technique^43^ in unconfined boundary conditions. To obtain the averaged stress map for average isotropic stress around defects, the stresses around each defect was rotated (head to tail axis for +1/2 half) along the defect and averaged as described in the figure legend.

### Velocity analysis

The velocities of the cells were obtained through particle image velocimetry (PIV) analysis using PIVlab in MATLAB. An interrogation window of size (64 x 64) pixels and (32 x 32) pixels, with an overlap of 50%, was used for the analysis. Having identified the location of defects with method described earlier, we obtained the average flow field around the defects identified by realigning these defects in a particular fashion. For +1/2 defect, we chose positive y-axis as head to tail axis and all the velocity values are rotated along this axis to obtain average velocity field.

### Average isotropic stress around fusion events

To characterize the average isotropic stress around fusion events, we took a square (50pixel x 50pixel, 32,5 µm x 32,5 µm) around each fusion event and isotropic stress was averaged within this square. Similarly, to obtain the isotropic stress around the fusion events in time, a square (50pixel x 50pixel) around each fusion event was average over each time point.

### Analysis of nuclear and fusion density

To get nuclear density around the defects or fusion locations, a square of area of (300 x 300 pixels, 180 x 180 µm) is drawn around it and the number of nuclei was counted within this square. Further, the number density is normalized with the global density in the movie area as [n/a]/[N/A] where ‘n’ is the number of nuclei within the square, ‘a’ is the area of the square, ‘N’ is number of nuclei globally and ‘A’ is the global area, respectively. Then, the normalized nuclear density is obtained over time for each defect or fusion locations. For this analysis, we specifically choose only the beginning of the movie to avoid overcounting the multinucleated cells.

To obtain fusion density around defects, with known fusion locations, the fusion coordinates were rotated along the defects within a given square and fusion density was estimated around the defects with hist3 function of MATLAB.

### Statistical correlation of topological defects with fusion events (cumulative probability)

To estimate the probability of correlation with defects, we estimated the cumulative probability of nearest defect from the fusion events. For each fusion event, the distance to its closest topological defects (+1/2 and -1/2 separately) was measured in the two preceding frames. We then computed the cumulative areal probability described previously^14^ using the formula P(r)=∑H(Δr/2−|r−ri|)/(2πrΔr) where H is Heaviside step function, and r ranges from r = 50 µm to r = 500 µm spaced by Δr = 50 µm. Furthermore, to distinguish the areal probability from random events, a set of random points (equal to fusion events in each frame) was uniformly generated.

### Statistical correlation of topological defects with fusion events (raw probability)

For each fusion event, the distance to its closest topological defects (+1/2 and -1/2 separately) was measured in the two preceding frames. We then computed the raw probability density (normalized) vs distance to closest defect for fusion events and random locations. We then estimate the kernel density using MATLAB built-in function. A significance test was done on the raw data in MATLAB with two sample KS test.

### Correlation of stress and fusion occurrence

For correlation analysis stable defects whose positions persist over long periods (≥ 12-16 hours) were chosen. For these stable defects, (i) the temporal evolution of the compressive stress magnitude and (ii) the number of fusion events occurring in their vicinity over the same interval was obtained. The temporal correlation of evolution of stress around stable topological defects and fusion occurrence with their pooled correlation was obtained in MATLAB.

### Continuum model of myotube monolayers

To model dynamics of myotubes, we use a dry model of compressible active nematics^46^. We describe nematic cell orientational order via order parameter, 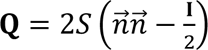, where *S* is the magnitude of the order, *n⃗* is the orientation of the cell order, and **I** is the identity tensor. Additionally, we introduce fibronectin orientation parameter, 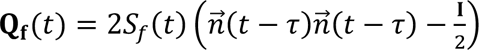, where, *τ* represents the time delay over which the orientation of fibronectin follows the orientation of the cells. Moreover, we assume that *S_f_*(*t*) is the monotonically increasing order parameter from 0 to 1, to model the increase in production of fibronectin over time:

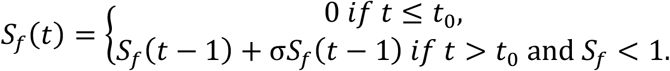

The dynamics of the cell nematic tensor is governed by

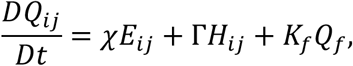

where co-rotational derivative is given by 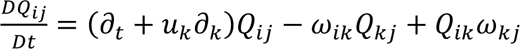 with *ω_ij_* = (*∂_j_u_i_* – *∂_i_u_j_*)/2 and *E_ij_* = (*∂_i_u_j_* + *∂_j_u_i_*)/2. Here, *χ* is the aligning parameter, Γ is the rotational diffusivity, and *H_ij_* is the molecular field. The last term in the equation denotes the alignment of nematic director to the fibronectin director.

The free energy given by Landau-de Gennes term and a 3-elastic constant terms^61^:

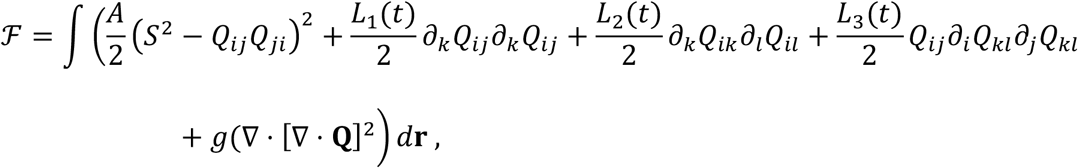

where

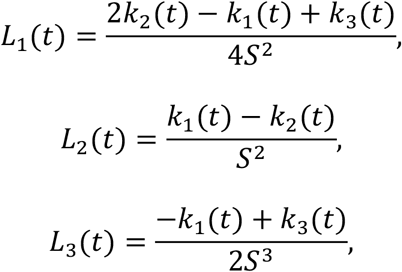

are time-dependent splay, twist and bend elastic constants from Frank elastic energy. In order to incorporate time-dependent increase of orientational elasticity due to cytoskeleton maturation, three elastic coefficients are given by

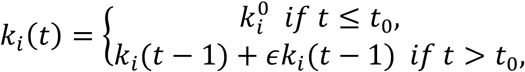

where *t*_0_ is the time at which the orientational elasticity starts to increase and is a positive coefficient. To avoid numerical instability, three-elastic constants are capped at maximal value, 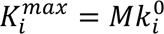. The last term in the free energy, ℱ, is the regularization term for numerical stability of the solutions. The first term is the nematic alignment term.

Since we are in the over-damped regime, the velocity is governed by the force-balance equation balancing the frictional damping with the divergence of the stress:

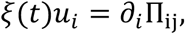

where *ξ*(*t*) is the time-dependent isotropic friction coefficient given by

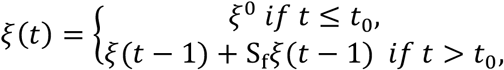

and **Π** = **Π**^elastic^ + **Π**^active^ is the stress tensor. Here,

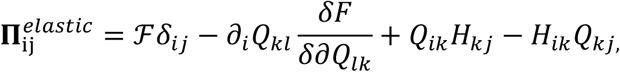

and

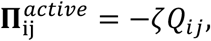

where *ζ* < 0 is the contractile activity.

We solve the equations numerically over a square 2D domain of size *L_x_* by *L_x_* with periodic boundary conditions. The initial three elastic constants are chosen such that the compressive stress is located at the head of the defect. We choose the maximal value of increase in orientational elasticity, *M*, such that it captures 3-fold decrease in the number of defects, observed in experiments. The parameter values used in the simulations are- rotational diffusivity (*Γ*)=0.1, initial splay elastic constant 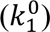=0.009, initial twist elastic constant 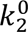 = 0.009, initial bend elastic constant 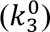=0.045, alignment parameter (χ)=0, dipolar activity (ζ)=-0.08, isotropic friction(ξ)=5, coupling constant (*A)*=0.1, 4th derivative term for num. stability(*g)=*0.1, orientational elasticity growth rate (ɛ)=0.0005, maximal increase in orient. Elasticity (*M*)=5, time when elasticity starts to increase (*t*_0_) =10500, size of computational domain (*L_x_*)=64, alignment to fibronectin (*K_f_*)=0.001, rate of increase of fibronectin order (σ)=0.0025, time-delay in fibronectin order (τ)=500, Domain size (*L_x_*)=64.

### Elastic constant approximation within nematohydrodynamics

By going beyond the one-constant approximation and explicitly distinguishing bend, splay, and twist elasticities within nematohydrodynamics, our model shows analytically how purely passive elastic anisotropy reshapes the isotropic stress field of defects. Specifically, when bend elasticity dominates, the head of a +1/2 defect is tensile; when splay elasticity dominates, the head becomes compressive—precisely as observed experimentally (Fig. 3e). In other words, without invoking changes in activity or substrate coupling, a shift in the relative orientational elasticities alone is sufficient to flip the defect stress polarity and amplify the magnitude of stress localization. According to theoretical studies of passive nematics^62^, the differences in the elastic response to orientational deformations affect the stress patterns of topological defects and we expect this to translate to active systems. By distinguishing bend, splay and twist deformation effects on orientational elasticity via introduction of 3-constant elasticity approximation^61^, we can use theory of nematohydrodynamics to analytically predict the stress patterns around defects for different values of elastic constants.

We define director of a +1/2 defect as

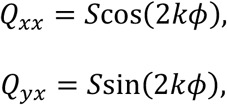

where *k* = 0.5 is the charge of the defect. Only elastic terms contribute to isotropic stress, given by

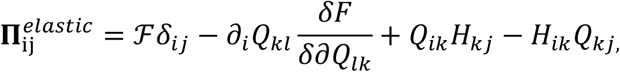

When bend constant dominates, the defect head is tensile. On the other hand, when splay constant dominates, we observe compressive head, reproducing the experimental pattern (Fig 3e).

Hence, we show how by changing only passive property of active material we can flip the stress pattern around the defects and increase the magnitude of compression/tension around the defect head (Extended Data Fig. 9a). In our model, the “theoretical stress” corresponds to the isotropic stress component that BISM infers from traction forces and these stress patterns are consistent with stresses measured via BISM.

## Acknowledgments

We thank the members of the “Cell Adhesion and Mechanics” and “Muscle Development, Growth and Repair” teams for helpful discussions. We acknowledge the ImagoSeine core facility of Institute Jacques Monod, member of France-BioImaging (ANR-10-INBS-04) and IBiSA, with support of Labex “Who Am I”, as well as the Centre d’Imagerie Quantitative Lyon-Est for imaging support. We thank Philippe Marcq for help with BISM. We thank Gerome Gros (Institut Pasteur) for the kind gift of the LifeActNeonGreen-IRES-tdTomatoMyl9 DNA construct.

## Author contributions

Conceptualization: YLT, SD, AA, AD, CM, BL; Methodology: YLT, SD, AA, VM, LB, AD, CM, BL; Investigation: YLT, SD, AA; Funding acquisition: AD, CM, BL; Project administration: CM, BL; Supervision: ED, AD, CM, BL; Writing – original draft: YLT, SD, AA, AD, CM, BL; Writing – review & editing: YLT, SD, AA, ED, LB, AD, CM, BL.

## Data and materials availability

Data, materials and image analysis code are available upon request.

