## Extended Data for "Extracellular matrix mediated stresses spatially bias myoblast fusion and myotube growth"

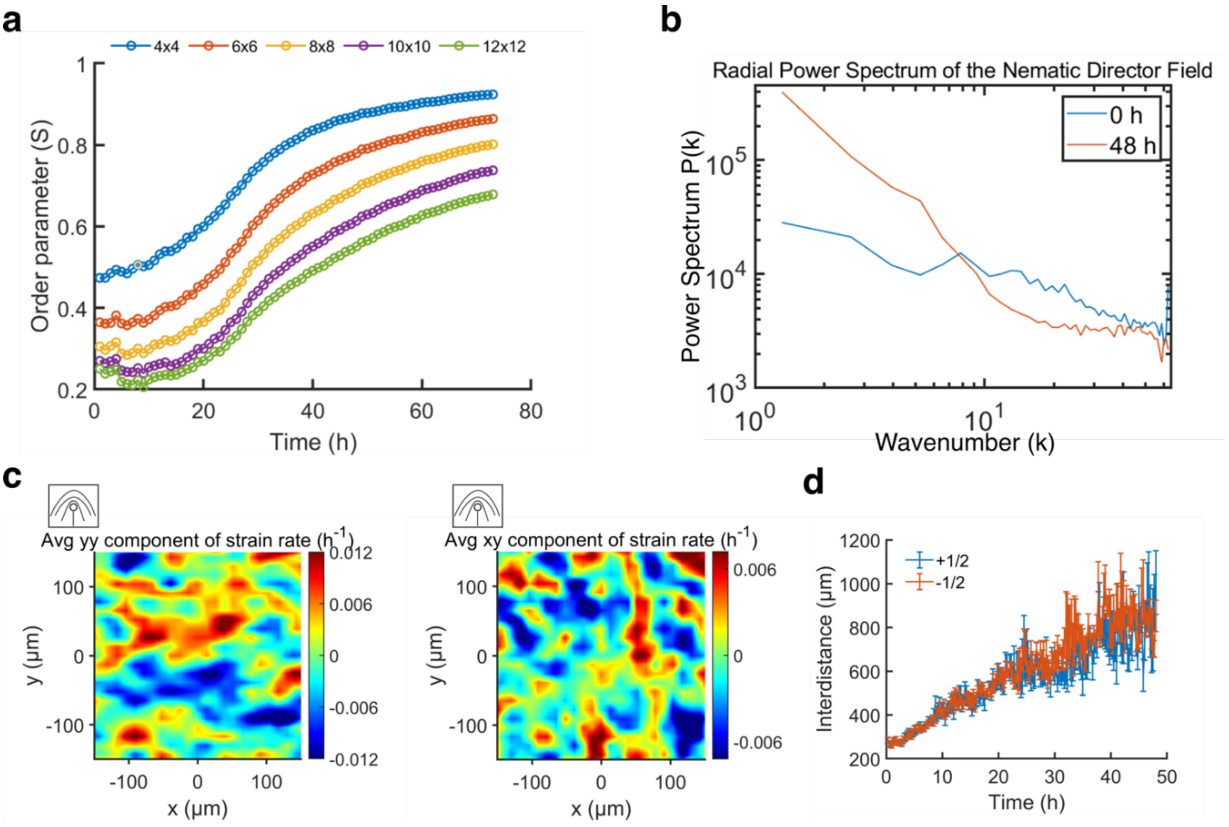


**Extended Data Fig. 1: Myoblasts display active nematic behavior and ordering at large scale during culture time**. a, Order parameter obtained at different window size of 4x4 -12x12 nematic directors (each director is of 90 pixels with a spacing of 45 pixels). b, Evolution of power spectrum with wave number at 0h and 48h time points (right). c, Strain-rate map around +1/2 defects showing yy-strain-rate (left) and xy-strain rate (right) component from around +1/2 defects (n=7503 defects from n=3 independent movies). Positive strain rate for yy-component at the head of +1/2 shows a typical of contractile system. d, Evolution of the interdistance between each type of topological defect along a 48h culture period (n=3 movies, error bars are S.E. from the mean).

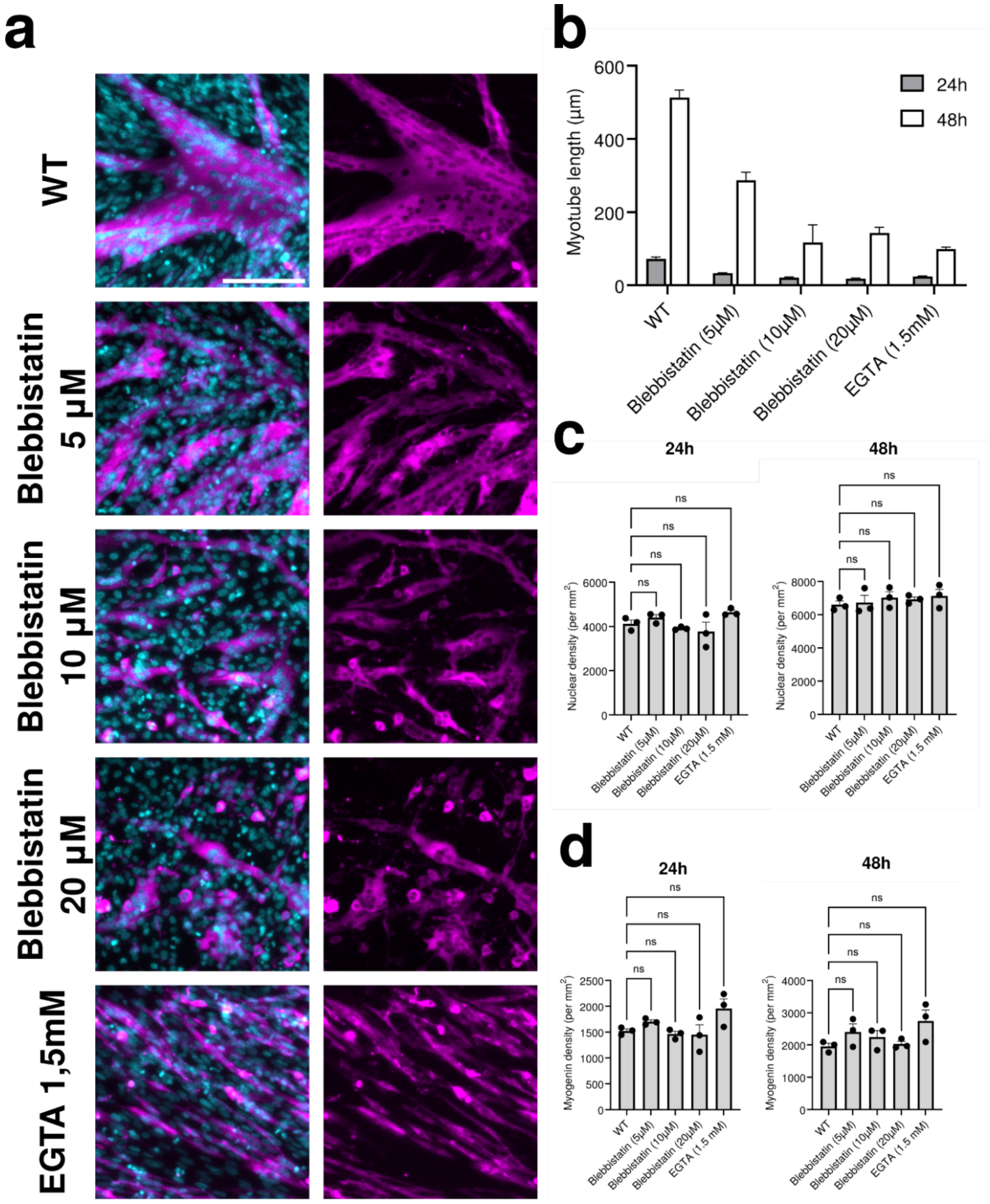
a
WT
Blebbistatin 5 µM
Blebbistatin 10 µM
Blebbistatin 20 µM
EGTA 1,5mM
b
Myotube length (µm)
600
400
200
0
24h
48h
WT
Blebbistatin (5µM)
Blebbistatin (10µM)
Blebbistatin (20µM)
EGTA (1.5mM)
c
24h
48h
ns
Nuclear density (per mm$^2$)
6000
4000
2000
0
8000
6000
4000
2000
0
WT
Blebbistatin (5µM)
Blebbistatin (10µM)
Blebbistatin (20µM)
EGTA (1.5 mM)
d
24h
48h
ns
Myogenin density (per mm$^2$)
2500
2000
1500
1000
500
0
4000
3000
2000
1000
0
WT
Blebbistatin (5µM)
Blebbistatin (10µM)
Blebbistatin (20µM)
EGTA (1.5 mM)

**Extended Data Fig. 2: Effects of Drug treatment on myotube growth and self-organization process**. a, Nuclei (cyan), MyHC (magenta) (left) and MyHC (right) staining of primary chicken myoblasts and myotubes fixed at 48h post induction of differentiation and treated with blebbistatin (5, 10, 20 μM), EGTA (1.5mM), or DMSO as control (WT). Scale bar, 200 μm. b, Average number of myotubes length at 24h and 48h time points (n=3 independent experiments). c, d Nuclear (c) and Myogenin positive nuclei (d) density at 24h and 48h time points for control and each drug treatment (n=3 independent experiments). Statistical comparisons were performed using one-way ANOVA to compare each treatment with the WT control. Statistical significance was defined as $p<0.05$. Error bars are S.E. from the mean.

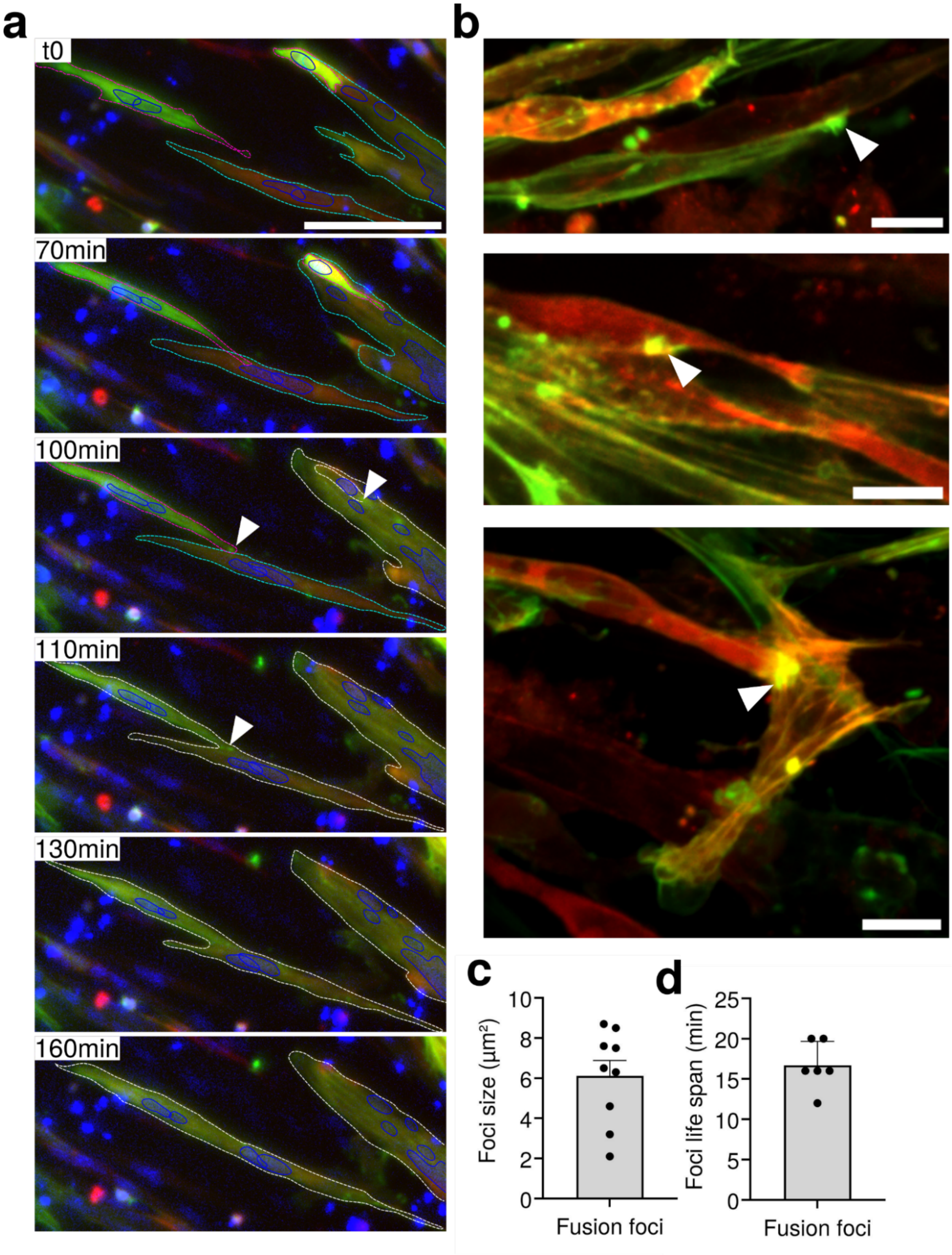


**Extended Data Fig. 3: Characterization of fusion events from foci detection to complete cell fusion**. a, Snap-shots of fusion events during the time-lapse. White arrowheads indicate the site of fusion characterized by an actin focus preceding cytoskeletal merging and fluorescence mixing; cells are delimitated by colored dotted lines (cyan, magenta) prior to the illustrated fusion events and white dotted lines once fusion occurred; actin (green), nuclei (blue) and myosin-II (red). Scale bar, 100 µm. b, High resolution snap-shots of fusion events illustrating actin foci at the contact site between cells with different orientation within the cell population. Scale bar, 10 µm. c-d, Quantification of actin foci size (n=9) and life span (n=6). Error bars are S.E. from the mean.

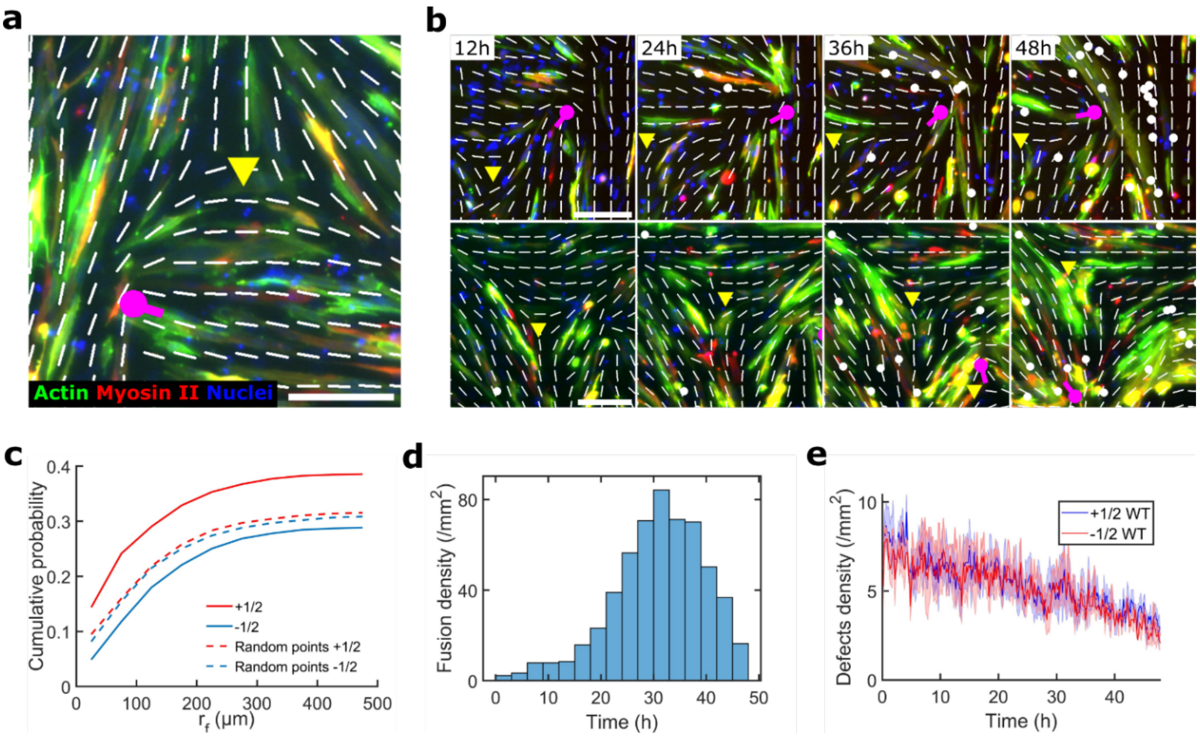


**Extended Data Fig. 4: Detection of the orientation field and topological defects and comparison with fusion localization**. a-b, Snapshots illustrating the detection of the orientation field (white lines) and the topological defects (+1/2, magenta dots and -1/2, yellow triangles), as well as the coordinates of fusion events (white dots), overlaying the fluorescent actin and myosin-II channels used for their detection. Scale bar, 200 µm. c, Cumulative probability to observe fusion events around topological defects in comparison to random points (n = 987 fusion events and random points, 8327 +1/2 defects and 7916 -1/2 defects over 3 independent movies). d, Evolution of the detected fusion density over the time course of the movies (n = 987 fusion events over 3 independent movies). e, Time evolution of the defect's density along the time course of the 3 independent movies used for fusion counting (error bars are S.E. from the mean).

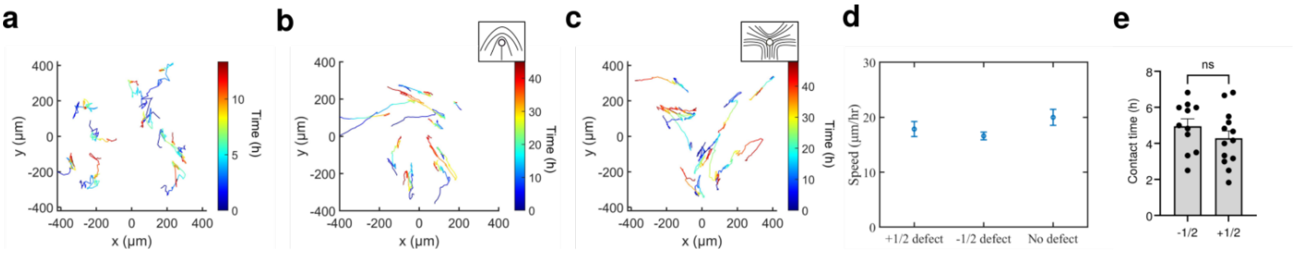


**Extended Data Fig. 5: Cell motility follows the organization of the cell population.** Examples of cell trajectories a before cell confluence (a), around a +1/2 defect (b) or -1/2 defect (c). Tracks are colored over time. d, Speed of cells around and far away from nematic defects (n=10 for each, error bars are S.E. from the mean). e, Cell-cell contact duration for plus half and minus half defects. Statistical significance was assessed using a Mann–Whitney test with $p = 0.25$ (error bars are S.E. from the mean).

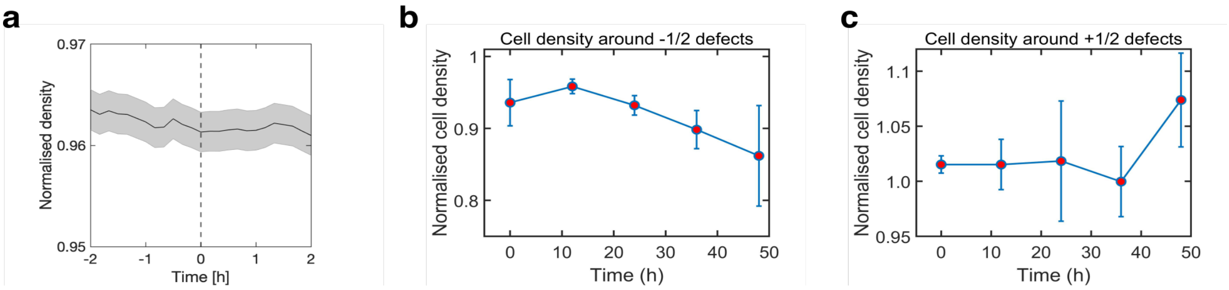


**Extended Data Fig. 6: Topological defects are not influenced by cell density**. a, Averaged normalized evolution of the cell density around the core of -1/2 defects. t0 corresponds to the birth of the defect respectively (analysis was performed over the first 24h of 3 independent 48h movies, error bars are S.E. from the mean). b-c, Averaged normalized cell density around the core of -1/2 (b) and +1/2 (c) defects present at specific time points during the culture period (3 independent movies, error bars are S.E. from the mean).

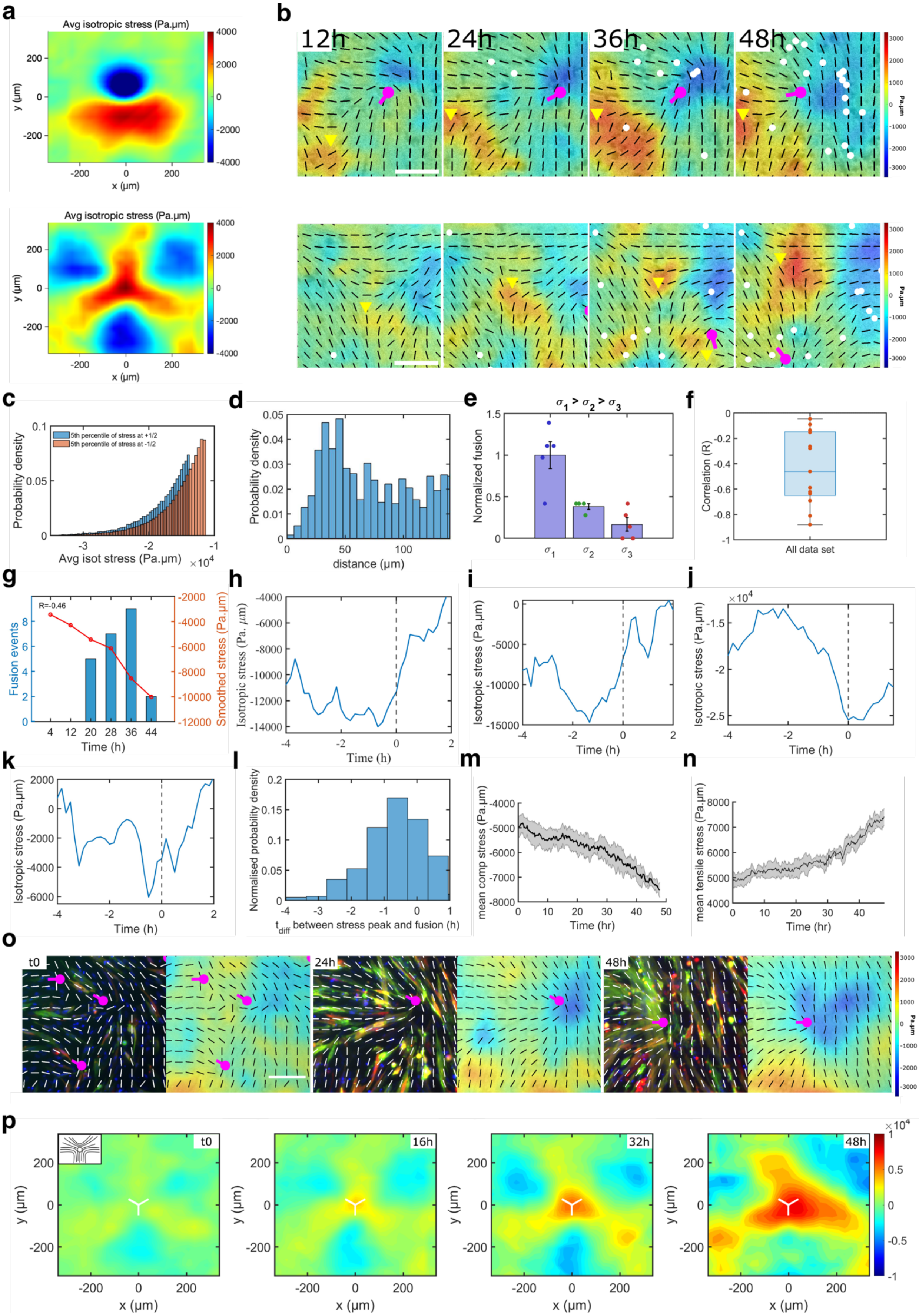

a
Avg isotropic stress (Pa.µm)
y (µm)
x (µm)
b
12h
24h
36h
48h
Pa.µm
c
Probability density
5th percentile of stress at +1/2
5th percentile of stress at -1/2
Avg isot stress (Pa.µm)
$\times 10^4$
d
Probability density
distance (µm)
e
$\sigma_1 > \sigma_2 > \sigma_3$
Normalized fusion
$\sigma_1$
$\sigma_2$
$\sigma_3$
f
Correlation (R)
All data set
g
R=-0.46
Fusion events
Smoothed stress (Pa.µm)
Time (h)
h
Isotropic stress (Pa. µm)
Time (h)
i
Isotropic stress (Pa.µm)
Time (h)
j
$\times 10^4$
Isotropic stress (Pa.µm)
Time (h)
k
Isotropic stress (Pa.µm)
Time (h)
l
Normalised probability density
$t_{diff}$ between stress peak and fusion (h)
m
mean comp stress (Pa.µm)
Time (hr)
n
mean tensile stress (Pa.µm)
Time (hr)
o
t0
24h
48h
Pa.µm
p
t0
16h
32h
48h
y (µm)
x (µm)
$\times 10^4$

**Extended Data Fig. 7: Evolution of isotropic stress around topological defects and correlation to fusion events.** a, Averaged isotropic stress map around +1/2 (top) and -1/2 (bottom) defects (n = 8327 +1/2 defects and n = 7890 -1/2 defects in 3 independent movies). b, Time projection of the coordinates of the fusion events (white dots) that took place in proximity to a +1/2 defect (top row, magenta) or -1/2 defect (bottom row, yellow triangle) during a 48h movie. White lines illustrate the averaged orientation of the cells overlaying the map the isotropic stresses taking place in these regions of the cell culture. Scale bar, 100 µm. c, 5$^{th}$ percentile of isotropic stress around +1/2 and -1/2 defects depicting higher compressive stress intensities around +1/2 defects d, Probability density of finding maximal compressive stress (higher than average compressive stress) around +1/2 defects showing higher compressive stresses near the core (<50 µm) of the defect. e, Fusion events enrichment at stable topological defects that associated with higher compressive stress (from 3 independent movies, error bars are S.E. from the mean). f, Correlation of compressive stress around stable topological defects with of the fusion events (n=13 defects from 3 independent movies, with mean R=-0.49 and pooled correlation significance of p=0.0016). g, An example of compressive stress evolution and fusion occurrence around stable defect. h-k, Snaps showing evolution of isotropic stress during fusion events for multiple events. l, Probability of time difference between stress peak and fusion occurrence (n=353 fusion events from 3 independent movies). m-n, Time evolution of the averaged compressive (e) and tensile (f) stress taking place in the 3 independent movies analyzed (error bars are S.E. from the mean). o, Time lapse illustrating the evolution of the myotubes (left, green-actin, red-myosin-II, blue-nuclei) and isotropic stress (right) around a +1/2 defect along the culture period. Scale bar, 100 µm. p, Evolution of average isotropic stress maps obtained for -1/2 defects present at each analyzed time point in the 3 movies analyzed. Inset is a schematic of the orientation field characteristic of +1/2 defects and central mark represents the core of the defect.

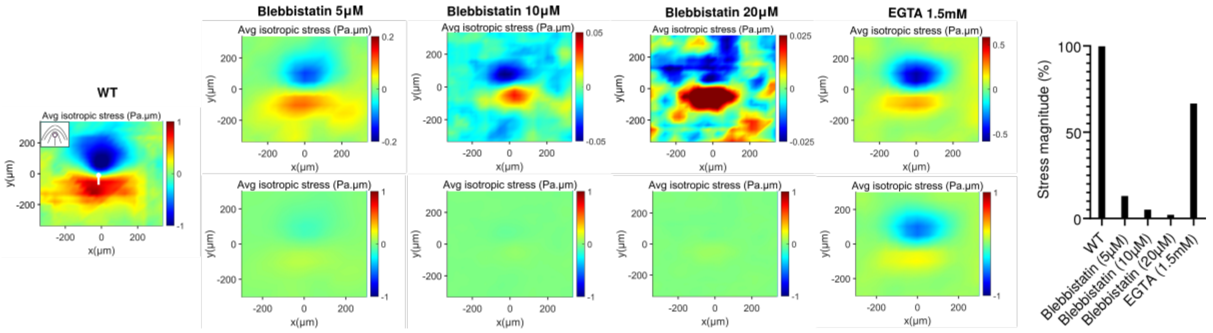


**Extended Data Fig. 8: Averaged stress map for various drug treatments**. Averaged isotropic stress around +1/2 half defects for sustained blebbistatin (5, 10, and 20µM), EGTA (1.5mM) and control DMSO treated cells (WT) (averaged over 3 independent movies and n>=2000 defects) (left). The bottom line shows the stress maps with a scale set on the control condition while the top row show the same maps with an adapted scale to illustrate the stress distribution preservation. Quantification of stress magnitude for sustained Blebbistatin (5, 10, 20µM) and EGTA (1.5mM) with respect to control (right).

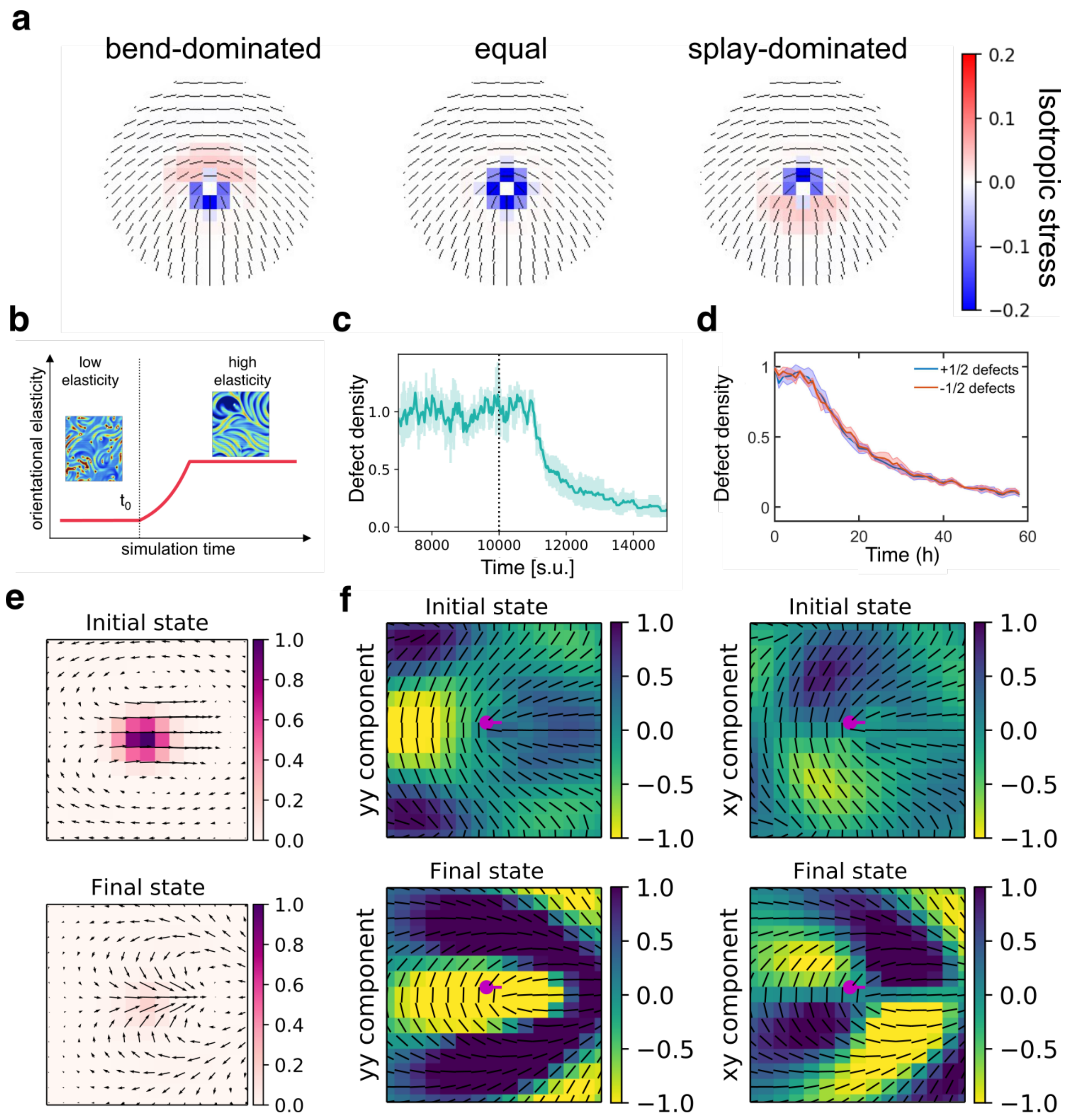


**Extended Data Fig. 9: Increase in orientational elasticity dictates stress pattern**. a, Analytically predicted isotropic stress patterns around defects in various regimes: bend-dominated, equal, and splay-dominated). b, Schematic depiction of the simulation setup of the active nematic with orientational elasticity (AN+OE). Once the system reaches active turbulence at $t_0$, orientational elasticity (3-constant approximation) is monotonically increased with time until saturating at a maximal value. This leads to visible changes in the isotropic stress patterns (inset plots above). c-d, Defect density obtained from AN+OE simulations (scaled by low-elasticity regime value) as a function of time for model (c) and experimental data (d, normalized, error bars are S.E. from the mean). e, Flow profiles around +1/2 defects for initial (top) and final states (bottom) for the simulations incorporating both orientational elasticity increase and coupling to ECM deposition (AN+OE+ECM). f, Strain rate yy (left) and xy (right) around +1/2 defects for initial (top) and final (bottom) states during simulations for the simulations incorporating both orientational elasticity increase and coupling to ECM deposition (AN+OE+ECM).

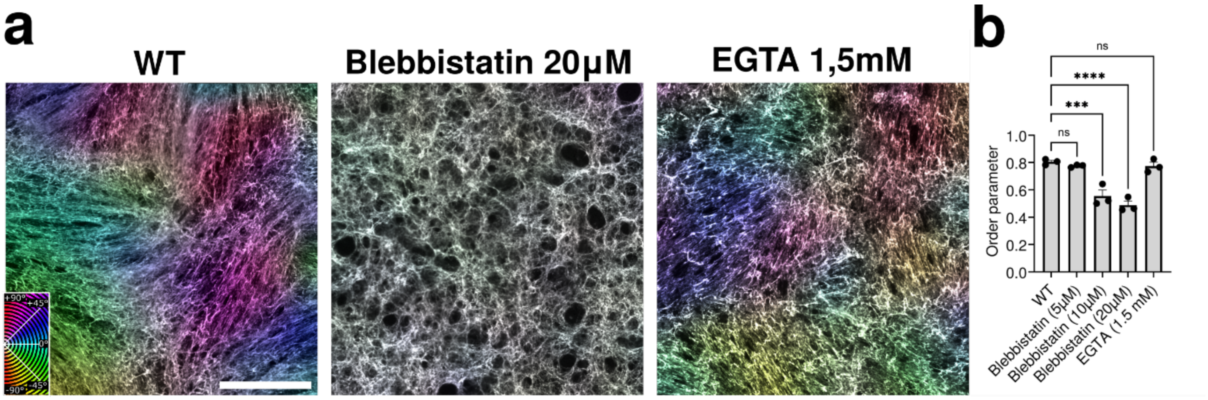


**Extended Data Fig. 10: ECM remodeling after drug treatment**. a, Snapshot showing fibronectin staining in control, Blebbistatin (20µM) and EGTA (1.5mM), pseudo-colored according to local orientation of fibronectin. Inset shows the color wheel applied to the local orientation of the cells. b, Order parameter of fibronectin ordering in control, blebbistatin (5, 10, 20µM) and EGTA (1.5mM). Statistical comparisons were performed using one-way ANOVA to compare each treatment with the WT control (n=3 independent experiments). Statistical significance was defined as $p<0.05$. Scale bar, 500 µm.
