## Supplementary Information for "Extracellular matrix mediated stresses spatially bias myoblast fusion and myotube growth"

**The PDF file includes:** Supplementary information contains Supplementary Fig.1-3 and supplementary video 1-9.

**Supplementary Information:**

Supplementary information contains Supplementary Fig.1-3 and supplementary video 1-9.

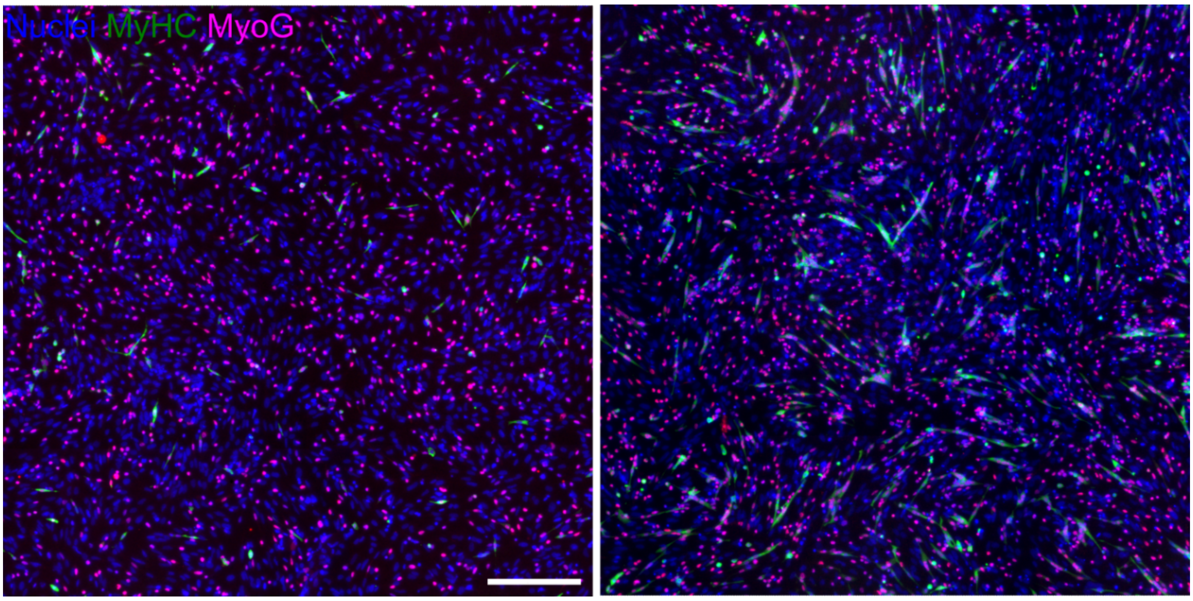


**Supplementary Fig. 1: Myogenin evolution during myotube growth and self-organization process.** Nuclei, MyHC, and MyoG staining of primary chicken myoblasts and myotubes fixed at 12h (left) or 24h (right). Scale bar, 500 µm.

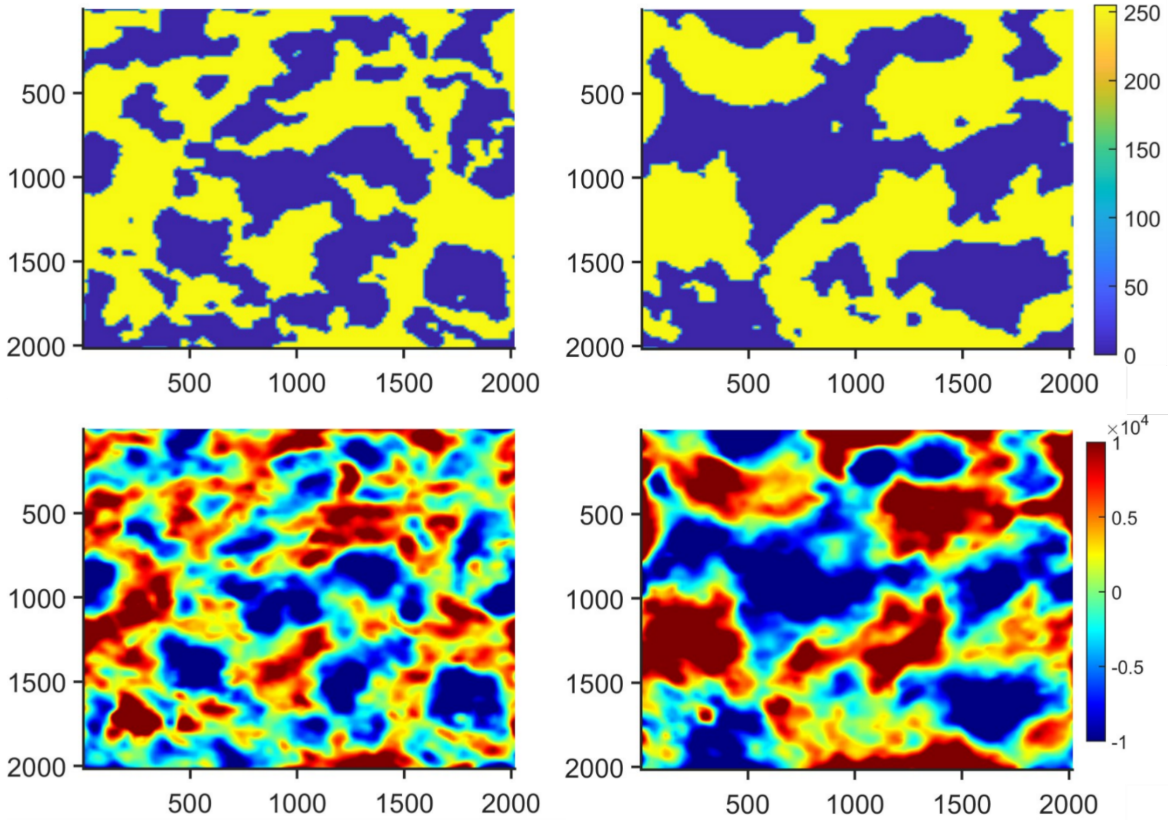


**Supplementary Fig. 2: Representative images showing evolution of compressive and tensile regions.** Increasing regions from beginning (left) to the end (right) of the movies using binary (0=blue, 255=yellow) maps (top) and isotropic stress (bottom).

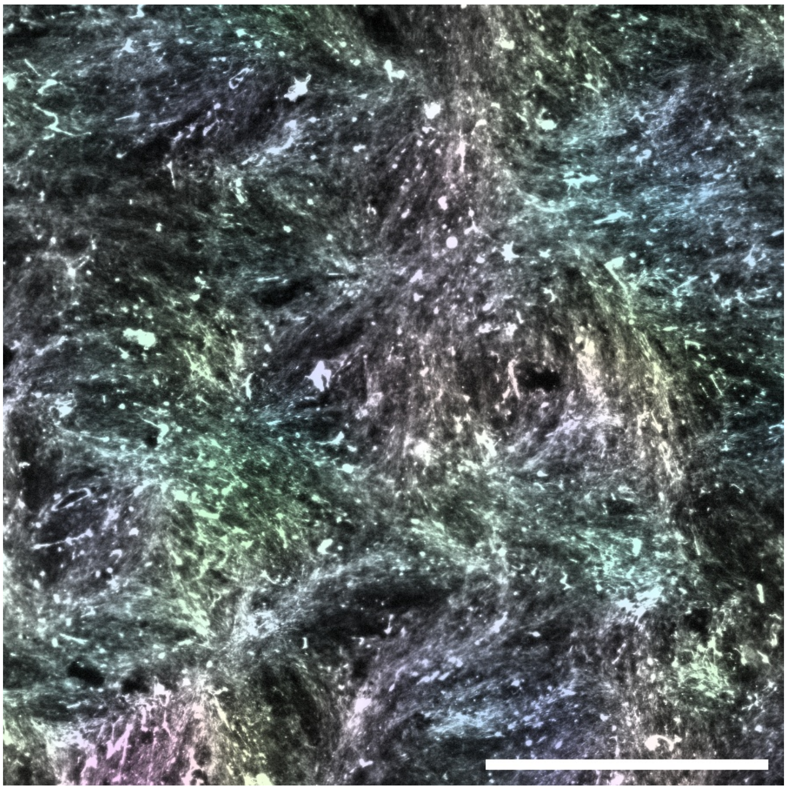

**Supplementary Fig. 3: ECM preservation after decellularization.** Snapshot showing fibronectin staining after decellularization, pseudo-colored according to local orientation of fibronectin. Scale bar, 500 µm.

**Supplementary Video 1: Chicken primary myoblasts transition to myotube in culture. Evolution of a primary chicken myoblasts culture along a 72h period. Scale bar, 500 µm.**

**Supplementary Video 2: Time lapse of a motile +1/2 defect from birth to annihilation.** Bright-field overlaid with averaged local orientation of the cells (red lines). +1/2 defects (magenta dots) and -1/2 defects (yellow triangle) merge together, leading to their annihilation. Scale bar, 100 µm.

**Supplementary Video 3: Effect of blebbistatin treatment on primary chicken myoblasts in culture.** Evolution of a primary chicken myoblasts culture along a 48h period. Scale bar, 500 μm.

**Supplementary Video 4: Evolution of the myoblast culture transduced with the lentiviral construct allowing to observe actin (green) and myosin-II (red) cytoskeleton, associated to Hoechst staining (blue) to observe the nuclei, along a 48h period.** Scale bar, 200 μm.

**Supplementary Video 5: Close-up area illustrating progressive myotube formation through fusion of myoblasts.** Actin (green) and myosin-II (red) and nuclei (blue). Scale bar, 100 μm.

**Supplementary Video 6: Close-up area illustrating fusion events occurring during myotube formation.** Actin (green) and myosin-II (red) and nuclei (blue). White arrowhead highlights actin foci preceding membrane merging and fluorescent signal mixing during the fusion process. Scale bar, 100 μm.

**Supplementary Video 7: Evolution of average isotropic stress maps (Pa. μm) obtained for +1/2 defects present at each analyzed time point in the 3 movies (48h) analyzed.**

**Supplementary Video 8: Evolution of average isotropic stress maps (Pa. μm) obtained for -1/2 defects present at each analyzed time point in the 3 movies (48h) analyzed.**

**Supplementary Video 9: Movie of single simulation realization showing the evolution of the active nematic system with coupling to ECM.** Right: isotropic stress (color bar) and director (black lines) of nematic. Left: director of fibronectin (blue lines). White and black circles denote +1/2 and -1/2 defects respectively.
